# Modelling Phenomenological Differences in Aetiologically Distinct Visual Hallucinations Using Deep Neural Networks

**DOI:** 10.1101/2023.02.13.528288

**Authors:** Keisuke Suzuki, Anil K. Seth, David J. Schwartzman

## Abstract

Visual hallucinations (VHs) are perceptions of objects or events in the absence of the sensory stimulation that would normally support such perceptions. Although all VHs share this core characteristic, there are substantial phenomenological differences between VHs that have different aetiologies, such as those arising from neurological conditions, visual loss, or psychedelic compounds. Here, we examine the potential mechanistic basis of these differences by leveraging recent advances in visualising the learned representations of a coupled classifier and generative deep neural network – an approach we call ‘computational (neuro)phenomenology’. Examining three aetiologically distinct populations in which VHs occur - neurological conditions (Parkinson’s Disease and Lewy Body Dementia), visual loss (Charles Bonnet Syndrome, CBS), and psychedelics - we identify three dimensions relevant to distinguishing these classes of VHs: realism (veridicality), dependence on sensory input (spontaneity), and complexity. By selectively tuning the parameters of the visualisation algorithm to reflect influence along each of these phenomenological dimensions we were able to generate ‘synthetic VHs’ that were characteristic of the VHs experienced by each aetiology. We verified the validity of this approach experimentally in two studies that examined the phenomenology of VHs in neurological and CBS patients, and in people with recent psychedelic experience. These studies confirmed the existence of phenomenological differences across these three dimensions between groups, and crucially, found that the appropriate synthetic VHs were representative of each group’s hallucinatory phenomenology. Together, our findings highlight the phenomenological diversity of VHs associated with distinct causal factors and demonstrate how a neural network model of visual phenomenology can successfully capture the distinctive visual characteristics of hallucinatory experience.

## 1 Introduction

Visual hallucinations (VHs) are perceptual experiences that occur in the absence of the sensory stimulation that would normally accompany such experiences. The decoupling of perceptual experience from sensation, together with the varied nature of VHs across different aetiologies, provides an opportunity to investigate the computational processes and neural mechanisms that may underlie them, with implications for those that may underlie visual experience in general.

VHs are commonly experienced as a result of certain neurological diseases, such as Parkinson’s disease (PD) (Barnes & David, 2001; Boubert & Barnes, 2015; Fénelon et al., 2000; Mosimann et al., 2006) and Lewy Body Dementia (LBD) (Harding et al., 2002). They may also occur as the result of marked visual impairment, as in Charles Bonnet Syndrome (CBS) (Abbott et al., 2007; ffytche, 2005; Nair et al., 2015; Schultz & Melzack, 1991; Teunisse et al., 1996; Yacoub & Ferrucci, 2011). VHs also occur in psychiatric conditions such as schizophrenia, although they are less frequent than auditory hallucinations (Bauer et al., 2011; Waters et al., 2014). VHs are also common features of psychedelic states, such as those produced by the ingestion of classical hallucinogens such as LSD, psilocybin and DMT (Nichols, 2016). The VHs associated with each of these aetiologies have distinctive phenomenological (experiential) characteristics, raising the question of what aspects of the underlying neurophysiology and dynamics are responsible for these differences.

A popular approach to understanding the basis of VHs is through the lens of ‘predictive processing’ (PP) theories of perception and brain function (Rao & Ballard, 1999; Friston 2005; Hohwy & Seth, 2020). These theories view perception as an iterative process in which the brain is always trying to optimise an evolving ‘best guess’ (Bayesian posterior belief) about the most likely causes of the sensory inputs it encounters (Clark, 2013; Knill & Pouget, 2004; Lee & Mumford, 2003). In standard PP, this is achieved by the reciprocal exchange of top-down perceptual predictions and bottom-up (sensory) prediction error signals in a continual process of prediction error minimisation, which approximates Bayesian inference on the causes of sensory signals. Importantly, the outcome of this (approximate) inference process depends on the relative influence of (top-down) predictions and (bottom-up) prediction errors, a balance which is mediated by the estimated precision (informally, the ‘reliability’) of sensory signals relative to perceptual predictions (Friston & Kiebel, 2009; Fletcher & Frith, 2009; Yuille & Kersten, 2006; Zarkali et al., 2019). Within this framework, VHs can be broadly understood as resulting from aberrant inference in which this balance is disrupted in some way (Friston 2005; Corlett et al., 2019; Powers et al., 2016; Zarkali et al., 2019).

Such aberrant inference can take many different forms. Some researchers have interpreted neurological VHs as resulting from overly strong perceptual predictions which overwhelm sensory prediction error signals (O’Callaghan et al., 2017; Powers et al., 2016). Others have focused on psychedelic VHs and adopted a hierarchical perspective, suggesting that psychedelic hallucinations occur due to a relaxation of high-level perceptual predictions (or Bayesian ‘priors’), which has the effect of increasing the influence of bottom-up signalling on perceptual inference (Alonso et al., 2015; Carhart-Harris & Friston, 2019; Swanson, 2018; Timmermann et al., 2018). However, despite accumulating empirical evidence linking perceptual experience to inference and prediction error minimization in both normal (Hardstone et al., 2021) and hallucinatory (Carhart-Harris & Friston 2019; Powers et al., 2016; Schartner & Timmermann, 2020) perception, the computational basis of phenomenological differences between different kinds of VH has remained unclear.

To shed light on this question, we take a computational neurophenomenology approach (Suzuki et al, 2017; Ramstead et al., 2022)]. Neurophenomenology emphasises the importance of capturing first-person descriptions of phenomena of interest (such as VHs) that are amenable to neurocognitive methodologies (Lutz 2002; Lutz et al. 2002; Gould et al. 2014). This approach becomes computational neurophenomenology when computational modelling is used to simulate specific properties of perceptual experience (rather than, for example, the functions associated with perception, such as classification or discrimination), and where the computational models used have useful interpretations with respect to theories of perception or neurophysiological mechanisms [see also phenomenological approaches in robotics e.g., Tani, J., (2016)].

### 1.1 Phenomenological variation in hallucinatory experience

While there are substantial phenomenological differences in VHs both across and within different aetiological categories, we focus on three dimensions which broadly characterise variations in VHs arising from neurological, CBS and psychedelic origins. These dimensions are complexity, veridicality, and spontaneity.

Complexity. VHs can generally be categorised as being either simple (e.g., shapes, flashes or grid-like lattice patterns) or complex (e.g., well-defined recognizable forms, such as objects or people). Neurological VHs are typically complex, featuring false perceptions of family, other people, or animals (Frucht & Bernsohn, 2002; Mosimann et al., 2006; Papapetropoulos et al., 2008). In contrast, the most commonly reported class of VH in CBS are simple (Abbott et al., 2007; ffytche & Howard, 1999; Santhouse et al., 2000), with complex VHs being reported less frequently (Schultz & Melzack, 1991; Teunisse et al., 1996). Psychedelic VHs also vary in their complexity. VHs arising from low doses are usually associated with visual distortion and/or simple VHs, such as brightly coloured geometric ‘form constants’ including lattices, cobwebs, tunnels and spirals (Díaz, 2010; Kluver, 1926; Nichols, 2016; Studerus et al., 2011), with complex VHs not as frequently reported (Kometer et al. 2011; Studerus et al. 2011; Schmid et al. 2015). At higher doses, complex VHs are more likely to occur, including fully formed VHs comprising visual scenes with elaborate structural content such as landscapes, cities, and galaxies, as well as specific forms including human figures and animals (Kometer et al., 2013; Liechti, 2017; Shanon, 2002; Strassman et al., 1994; Studerus et al., 2011). The rarity of complex psychedelic VHs of this sort may reflect either their relative rarity and/or the difficulties inherent in providing subjective reports in high dose situations (Kometer et al., 2013; Kometer & Vollenweider, 2018; Studerus et al., 2011).

Veridicality. Neurological and CBS complex VHs typically display a high degree of perceptual veridicality. That is, they are reported as being similar in visual quality to normal perceptual experience (Barnes & David, 2001; Boubert & Barnes, 2015; Mosimann et al., 2006; Schultz & Melzack, 1991; Teunisse et al., 1996). There are some exceptions: for example, complex VHs in CBS are sometimes described as distorted or cartoon-like, however, the majority are described as vivid and life-like (Abbott et al., 2007; ffytche, 2005; ffytche & Howard, 1999; Santhouse et al., 2000; Schultz & Melzack, 1991). In contrast, psychedelic complex VHs are typically reported to have a lower degree of veridicality, which may not be surprising thanks to their dream-like qualities (for a review, see Sanz et al., 2018) and the prominence of visual distortion, unrealistic colours, patterns and kaleidoscopic imagery (Carhart-Harris et al., 2016; Preller & Vollenweider, 2018; Studerus et al., 2011; Timmermann et al., 2018). Indeed, some have suggested that the alterations in visual perception induced by psychedelics, such as psilocybin, rarely represent true hallucinations because, at least at moderate doses, they can be readily distinguished from real perceptions (Preller & Vollenweider, 2018).

Spontaneity: Neurological and CBS complex VHs typically occur spontaneously, that is, they are not experienced as being transformations of existing content within a perceived scene (Frucht & Bernsohn, 2002; Mosimann et al., 2006; Papapetropoulos et al., 2008; Schultz & Melzack, 1991; Teunisse et al., 1996). Spontaneous VHs, therefore, correspond closely to the folk-psychological idea of hallucination as perception in the absence of sensory stimulation. In contrast, anecdotal reports of psychedelic complex VHs describe them as frequently being driven by visual ‘seeds’ from within a perceived scene, bearing similarity to the everyday phenomenon of ‘pareidolia’ – or ‘seeing patterns in noise’ (Kometer & Vollenweider, 2018; Preller & Vollenweider, 2018; Swanson, 2018).

From this short review, it is possible to broadly characterise each aetiological distinct category of VH in terms of complexity, veridicality, and spontaneity. Neurological VHs typically display high veridicality, are spontaneous, and are mainly complex in nature. CBS VHs also typically display high veridicality, and are also spontaneous, but can occur in both simple and complex forms (see Table 1). Psychedelic VHs generally display lower veridicality compared to Neurological and CBS VHs, they tend to not be spontaneous and can occur in both simple and complex forms. We used these phenomenological profiles to tune the parameters used to generate synthetic VHs representative of each category.

**Table 1.** Summary of phenomenological characteristics associated with neurological, CBS and psychedelic VHs (clear) and the corresponding model manipulations (shaded) used to simulate these characteristics. Veridicality refers to the similarity in the visual quality of a given experience compared to normal visual experience. AM Type refers to the use of either DGN-AM or DCNN-AM. Complexity refers to the occurrence of simple or complex visual experience. Layer of DCNN describes the level within the DCNN that AM terminates. Spontaneity refers to whether the visual experience is dependent or independent of sensory input. Type of Error refers to whether Fixed, Deep-Dream or WTA error function was used. To simulate CBS VHs, we manually blurred the centre of each input image to mimic the central visual deficit associated with CBS.

| Type of Visual Phenomenology | Veridicality | AM Type | Complexity | Layer of DCNN | Spontaneity | Type of Error Function |
| --- | --- | --- | --- | --- | --- | --- |
| Benchmark | High | DGN-AM | Complex | Highest layer ('fc8') | Dependent | WTA |
| Neurological Complex VH | High | DGN-AM | Complex | Highest layer ('fc8') | Independent | Fixed |
| CBS Complex VH | High | DGN-AM | Complex | Highest layer ('fc8') | Independent | Fixed |
| CBS Simple VH | High | DGN-AM | Simple | Lower layer ('conv4') | Independent | Fixed |
| Psychedelic Complex VH | Low | DCNN-AM | Simple | Lower layer ('conv4') | Dependent | Deep-Dream |
| Psychedelic Simple VH | Low | DCNN-AM | Complex | Highest layer ('fc8') | Dependent | Deep-Dream |

### 1.2 Simulating dimensions of hallucinatory phenomenology

To simulate these specific aspects of hallucinatory phenomenology, we used the coupled neural network architecture of Nguyen et al., (2016), which combines pre-trained classifier (DCNN) and generative (DGN) networks (with no additional training), to generate synthetic snapshots of aetiologically distinct VHs. Intuitively, within this architecture, the image fed into the model can be viewed as analogous to visual input and the synthetic image produced by the model, the resulting perceptual experience.

The class of DCNN used by Nguyen et al. (2016) is a pre-trained feedforward hierarchical neural network (taken from Krizhevsky et al., 2017) that was trained through supervised learning to perform object recognition on the natural image dataset, ImageNet (ILSVRC 2012, Krizhevsky et al., 2017; Szegedy et al., 2015). The DGN used by Nguyen et al., (2016) was pre-trained independently from the DCNN (DGN taken from Dosovitskiy and Brox 2016a), using a Generative Adversarial Network (GAN) approach, a learning architecture that is capable of generating high-dimensional photorealistic images (Goodwell 2014).

DCNNs trained for natural image recognition are highly complex, with many parameters and nodes, such that their analysis requires innovative visualisation methods. A popular solution to this challenge has been to use algorithms that visualise the information processed by a target neuron (or group of neurons) within a DCNN. Activation maximisation (AM) is one such visualisation technique, which works by iteratively calculating the error between the predicted value and the actual value through an error function to maximise the activity of a single target neuron within a DCNN. The errors are passed through the network via backpropagation to alter the colour of each pixel within an input image to maximise activity within the target neuron (Erhan et al., 2009; Mahendran & Vedaldi, 2014; Simonyan et al., 2014). This process changes the image rather than altering the network to match the features of the image with what is represented by the target neuron. Due to this form of AM only operating within the DCNN, we refer to it as DCNN-AM. A slight modification of this approach is the well-known Deep-Dream visualisation algorithm, which applies AM across a user-defined layer of the DCNN instead of a single target neuron (Mordvintsev et al., 2015).

#### Veridicality

Previous studies using DCNN-AM have shown that it typically produces unrealistic, uninterpretable images (Erhan et al., 2009; Mahendran & Vedaldi, 2014; Simonyan et al., 2014). This is due to the vast set of possible images that may excite a target neuron, making it likely that the image produced will not resemble the natural images that the neuron has learned to detect. To produce more ‘human-interpretable’ images, Nguyen et al., (2016) developed a modified version of AM that utilised the learned natural image prior of a pre-trained DGN, which they refer to as DGN-AM. For each iteration, DGN-AM uses the pre-trained DGN to synthesise a new output image that maximises the activity of a target neuron within the DCNN, instead of updating the input image directly as in DCNN-AM. Using this approach Nguyen et al., (2016) found that the natural image prior of the DGN produced output images that were much more realistic than those produced by standard DCNN-AM.

To simulate differences in the veridicality of aetiologically distinct VHs, we therefore generated synthetic VHs both with (high veridicality), and without (low veridicality), the natural image prior provided by the DGN.

#### Spontaneity

A key challenge when designing network visualisation techniques centres on the parameters of the error function used to select target neuron(s). DCNN-AM and DGN-AM use a Fixed error function that returns an error for a pre-selected (user-defined) target neuron, resulting in visualisations that are independent of the input image. In contrast, the Deep-Dream error function, used in our previous simulations of hallucinatory phenomenology (Suzuki et al. 2017), returns errors for all the neurons within a specific layer of the DCNN that were activated by the input image, resulting in visualisations that are dependent on the input image.

To simulate the reported differences in the VH spontaneity we therefore selectively altered the error function used to select the target neuron(s) within the DCNN. To simulate spontaneous VHs we used error functions that are not dependent on the input image (Fixed), and to simulate VHs that are driven by visual ‘seeds’ from within a perceived scene we used an error function that is dependent on the input image (Deep-Dream).

Another reason we selected this approach is that it relates to the theorised disruption between (top-down) predictions and (bottom-up) prediction errors, which has been proposed to underlie VHs within PP accounts of perception (Alonso et al., 2015; Carhart-Harris & Friston, 2019; Swanson, 2018; Timmermann et al., 2018). Within this framework, the use of a predetermined target neuron by the Fixed error function can be viewed as simulating the overly strong perceptual predictions thought to be the cause of neurological/CBS VHs (O’Callaghan et al., 2017; Powers et al., 2016). The Deep-Dream error function can be viewed as simulating the increased influence of bottom-up signalling on perceptual inference thought to underlie psychedelic VHs, because in this case the selected target neurons are dependent on, and driven by, the properties of the input image.

#### Complexity

To simulate differences in the complexity of hallucinatory content we applied both forms of AM in a layer-specific manner, reasoning that simple hallucinatory phenomenology could be simulated by restricting the layer that AM terminates to lower layers of the DCNN, resulting in an overemphasis of low-level visual features extracted by these layers during training. To simulate complex hallucinatory phenomenology, we instead restricted the layer that AM terminates to higher layers.

Finally, to provide a benchmark of the Nguyen et al., (2016) model’s performance against which the above simulations of hallucinatory phenomenology could be compared, we produced simulations of non-hallucinatory (veridical) perceptual phenomenology. Normal perceptual experience is naturally characterised by high veridicality and a close dependence between visual input and perceptual experience. To mimic these features, we used DGN-AM and developed a new class of error function, which we call Winner-Take-All (WTA). The WTA error function returns an error for the neuron that was maximally activated by an input image, resulting in a balance between sensory signals and perceptual predictions (See section 2.1.2). The idea of this benchmark is to create an image set using the same network architecture when it is not tuned to model a VH of any kind.

Using the above approach, we set out to test if the coupled DCNN-DGN neural network architecture of Nguyen, et al., (2016) could be used to simulate three aspects of neurological, CBS and psychedelic hallucinatory phenomenology: veridicality, spontaneity, and complexity. Summarising, we simulated these three phenomenological dimensions by the inclusion or omission of the natural image prior of the DGN (veridicality), altering the error function used to select the target neuron(s) within the DCNN (spontaneity), and restricting the level within the DCNN that AM terminates (complexity).

### 1.3 Do synthetic VHs provide an accurate representation of hallucinatory experience?

To draw conclusions about the validity of the synthetic VHs produced by this approach, it is important to establish the extent to which they match the subjective experience associated with each group’s hallucinatory experience. To investigate this question, we performed two additional experimental studies. In the first in-person study we developed a semi-structured interview that was used to enquire about the visual phenomenology associated with CBS and neurological VHs. The second online study recruited participants with recent psychedelic experience to assess how closely a recalled psychedelic experience matched example of the appropriate synthetic VHs. Both studies enquired about the general phenomenology, including specific questions regarding the complexity, veridicality, and spontaneity of their VHs and critically, also asked participants to directly rate the visual similarity between the appropriate synthetic VHs and their hallucinatory experiences.

## 2 Materials and Methods

### 2.1 DCNN-DGN model

Here we summarise the details of the coupled DCNN-DGN model architecture and DGN-AM visualisation algorithm used by Nguyen et al., (2016). Note that we used the original Nguyen et al., (2016) model architecture without any modifications. Modelling distinct classes of VH was instead achieved by applying the following modifications to the visualisation algorithm: 1) we added a function that allowed us to switch between the DGN-AM and DCNN-AM visualisation algorithms. 2) we added a function that allowed us to use different error functions when optimising images for both forms of AM. 3) We modified both forms of AM so that they could be applied in a layer-specific manner to the DCNN.

#### 2.1.1 Model Architecture

The DCNN used by Nguyen et al., (2016) is the CaffeNet architecture (Jin 2014), a minor variant of the AlexNet architecture (Krizhevsky et al., 2017). The DCNN was pre-trained for image classification based on the thousand categories of ImageNet, a dataset of natural photographs (Russakovsky et al., 2015). The CaffeNet architecture consists of five convolutional layers (c1-c5) and three fully connected layers (fc6, fc7, and fc8)(Figure 1). The layer fc8 is the highest layer of the DCNN (pre-softmax) and has 1000 outputs, each corresponding to one of the ImageNet categorical labels. During training, the visual characteristics of ImageNet photographs are extracted across all layers of the DCNN. The network learns via backpropagation to associate these features with distinct categorical labels. Consequently, the trained network implements a mapping from the pixel level of the input image to the respective categorical labels, represented as activation within specific neurons within the highest layer of the network (fc8).

**Figure 1.**
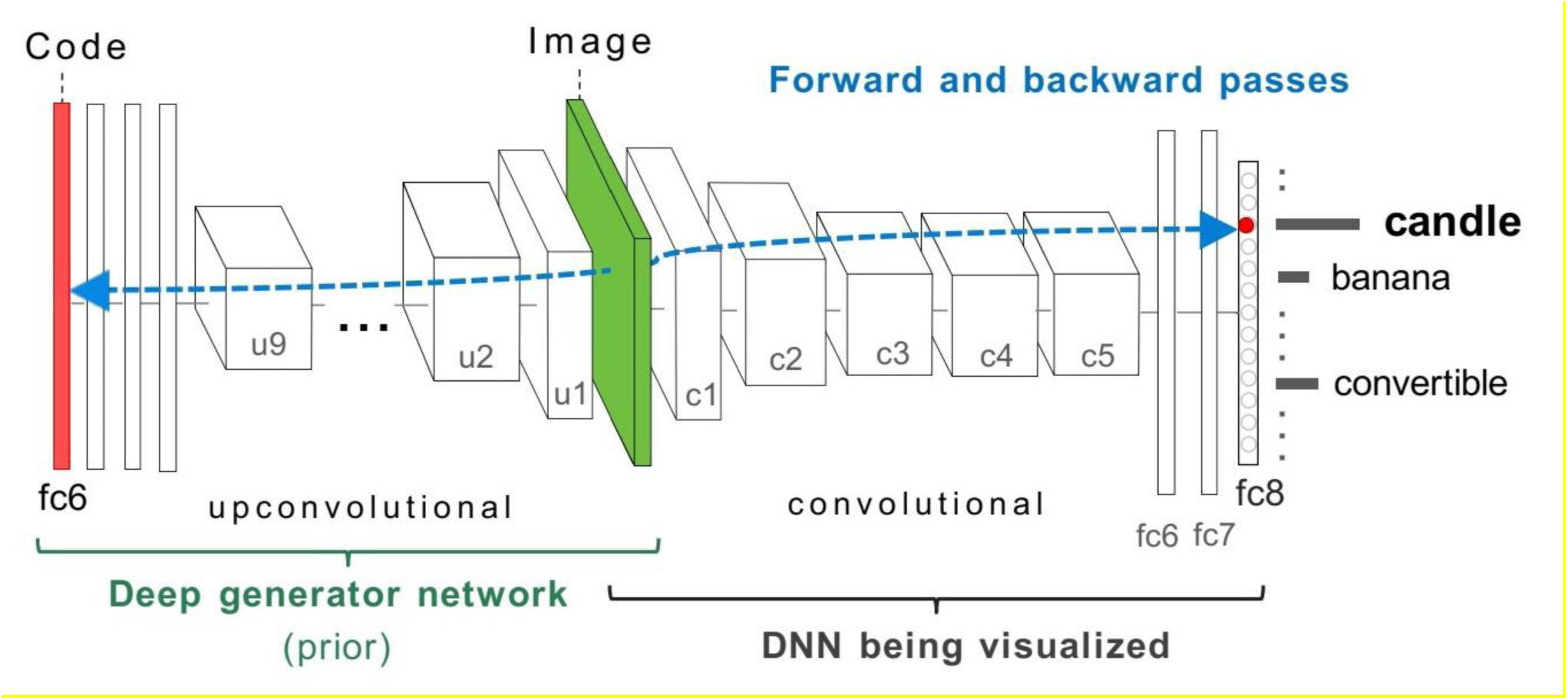
Nguyen et al., (2016) model architecture used in this study. The two networks, DCNN (labelled DNN) on the right, and DGN on the left are combined via the Image layer at the bottom of both networks (green). To synthesise a preferred input for a fixed target neuron (representing a “candle”) located in layer fc8 of the DCNN, the latent vector layer (red bar) of the DGN is optimised to produce an image that highly activates the target neuron. The gradient information (blue-dashed line) flows from the layer containing the target neuron in the DCNN via backpropagation through the image to the latent vector layer of the DGN. This process generates a new image that maximally activates the target neuron within the DCNN. Image adapted with permission from Nguyen et al., (2016).

The DGN used by Nguyen et al., (2016) was taken from Dosovitskiy & Brox, (2016a) and was pre-trained using the GAN framework, combined with the representation learning method (Figure 1)(see supplementary material for details of the GAN training). This DGN consists of nine up-convolutional (up-sampling and a subsequent convolutional) layers (u1-u9) and three fully connected layers (fc5, fc6, fc7) (Figure 1) which are designed to invert a convolutional network.

#### 2.1.2 Synthesising preferred images using AM

Here, we describe the two different classes of AM architecture that we used to generate synthetic VHs. The code used to generate all synthetic images is freely available here https://github.com/ksk-S/ModellingHallucinations

##### DCNN-AM

The core idea behind AM is simple: generate an input image that maximises the activation of a target neuron(s) of interest. Instead of updating the weights between the neurons in the network as occurs during training, AM iteratively modifies the image in a way that maximises the activity of the target neuron (see supplemental material for a formal description). This is done by defining an error function to return larger errors when the activation of the neuron is low and smaller errors when the activation is high. Note that AM inverts the relationship between the inputs and outputs of the DCNN, such that the network input is now the selected target neuron, and the output is the modified image.

##### DGN-AM

Because DCNN-AM by itself typically leads to unrealistic images, Nguyen et al., (2016) used a form of AM that combines a DGN with the DCNN (Figure 1), so that activity of the target neuron within the DCNN is maximised indirectly through the synthesis of a new image by the DGN (DGN-AM) (see supplemental material for a formal description). In DGN-AM, the gradient information does not terminate at the bottom layer of the DCNN but is passed through the image layer to optimise the latent vector of the DGN, which is then used to generate a preferred image for the selected target neuron (Figures 1 and 2).

**Figure 2.**
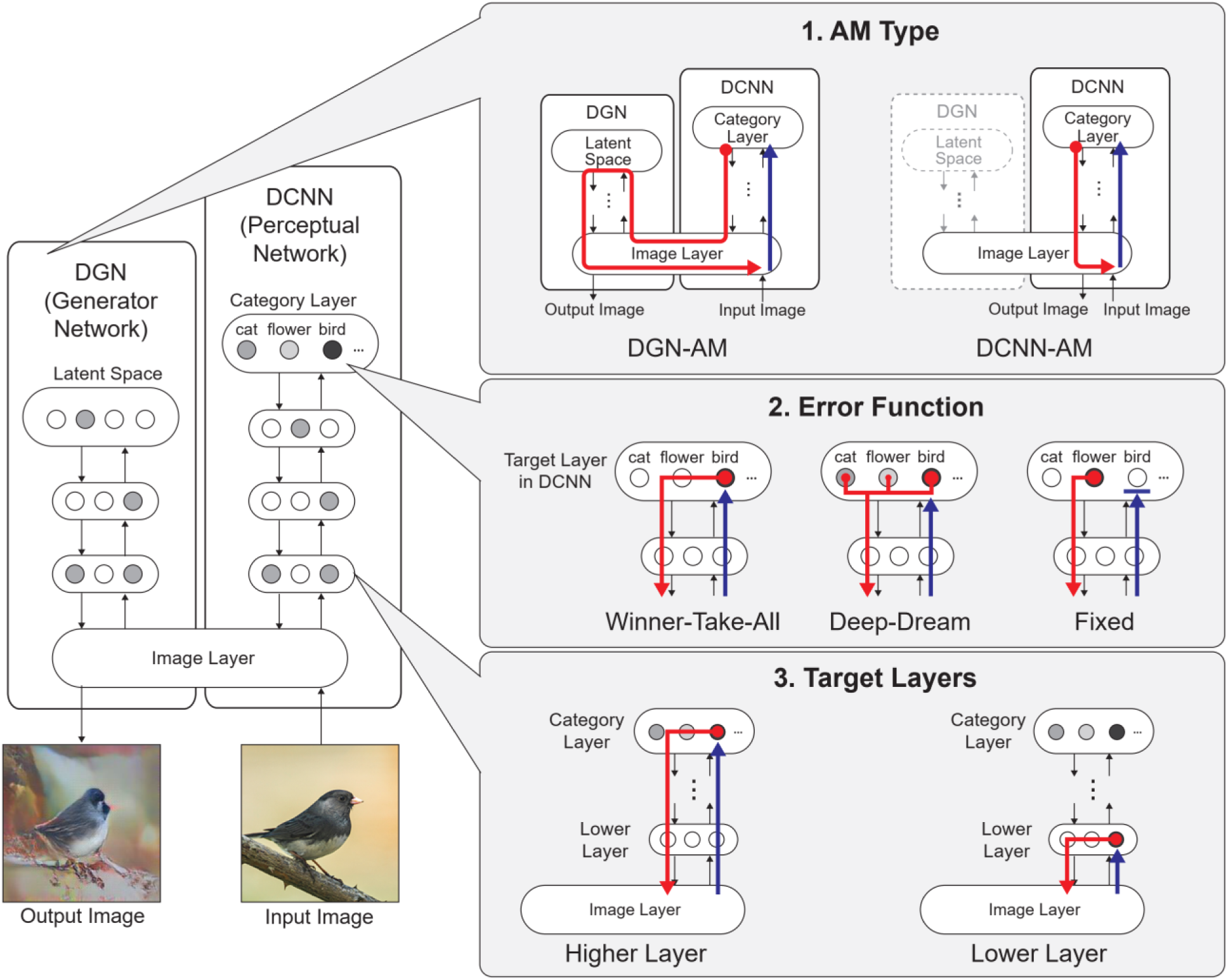
Left. Model architecture adapted from Figure 1. The model consists of a DGN (left) that constrains the network towards producing realistic images that maximally activate a target neuron within the DCNN (right). Right. The three independent manipulations that we applied to simulate specific forms of hallucinatory phenomenology. AM Type (top), we used two forms of AM, that either iterated through just the DCNN (DCNN-AM) or through both the DCNN and DGN (DGNN-AM). Error Function (middle) returns errors at the terminated layer for either a single predetermined neuron (Fixed), all neurons activated by the input image (Deep-Dream) or the single neuron that is maximally activated by the input image (Winner-Take-All). Target Layers (bottom) allows the selection of the specific layer that the AM process terminates at and returns errors from. Note that these three manipulations can be applied orthogonally.

#### 2.1.3 Modifying the parameters of AM: error functions and target layers

As previously mentioned, the error function implemented by AM is used to select the target neuron(s) to be optimised by the network. The application of different error functions has been shown to dramatically alter the output of the network (Simonyan et al., 2014; Mordvintsev et al 2015; Dosovitskiy A, Brox T. 2016b), a dependency we take advantage of to simulate the reported differences in the spontaneity of VHs.

##### Fixed error function

DCNN-AM and DGN-AM were both designed to visualise a single user-selected target neuron, meaning that the error function operates on the selected target neuron and returns errors that activate only this target neuron, while all other neurons return zero errors (Figure 2.1). Consequently, the output image is optimised to activate the selected target neuron only (usually representing a particular object category thanks to the DCNN/DGN training regime) regardless of the content of the input image.

##### Deep-Dream error function

In contrast to the above, in the Deep-Dream algorithm, the user selects which layer of the DCNN the error function is to operate on. Within the selected layer, multiple target neurons may be activated by the error function depending on which are activated by the visual features of the input image (Suzuki et al. 2017; Mordvintsev et al., 2015). The output image is therefore generated by a dynamic interplay between the visual features contained within the image and the neurons that are activated within the user-defined layer of the DCNN.

##### Winner-Take-All error function

In addition, we also use a new type of error function that combines the features of the Fixed and Deep-Dream approaches, which we call ‘Winner-Take-All’ (WTA). In WTA, as with Deep-Dream, the user selects which layer of the DCNN the error function is to operate so that the target neuron is selected by the visual features of the input image, however, only a single neuron with the highest activation to the input is selected as the target neuron, while the errors of all the other neurons are set to zero, as with Fixed. These three types of error functions are illustrated in Figure 2.2.

##### Target Layer

AM can be applied to any layer of the DCNN by terminating the upward pass at the specified target layer and beginning backpropagation from this layer (Mahendran & Vedaldi, 2014; Simonyan et al., 2014). In this study, we used either the top categorical layer (fc8) or lower layers (c3 and c4) of the DCNN (Figure 2.3).

Figure 2 summarises the manipulations to AM we used to simulate hallucinatory phenomenology from aetiologically distinct groups: AM type (DCNN-AM or DGN-AM), error functions (Fixed, Deep-Dream, or WTA), and user-selected target layer within the DCNN (Lower or Higher).

#### 2.1.4 Parameters used to simulate aetiologically distinct VHs

Based on the phenomenological profile of each aetiologically distinct category of VH (in terms of their complexity, veridicality, and spontaneity), we applied specific combinations of AM type, error function, and target layer that were designed to simulate each profile.

To do this, we first needed a benchmark, against which simulations of hallucinatory phenomenology could be compared. To provide this benchmark we used the model to simulate non-hallucinatory (veridical) perceptual phenomenology. We did this by applying DGN-AM with WTA and selecting the top layer of DCNN (fc8).

Next, to simulate variations in the veridicality of hallucinatory content we generated synthetic VHs using either DGN-AM (high veridicality) or DCNN-AM (low veridicality); i.e., with (DGN) or without (DCNN) including a natural image prior. We reasoned that because DGN-AM can produce realistic-looking, interpretable outputs it could be used to simulate the reported high veridicality of (complex and simple) neurological and CBS VHs (see Table 1)(Nguyen et al., 2016). In contrast, our previous work using DCNN-AM demonstrated its suitability to simulate the reduced veridicality of both simple and complex psychedelic VHs (Figure 2.1) (Suzuki et al. 2017).

To simulate differences in the reported spontaneity of complex VHs, we leveraged differences in error functions that are either dependent on (Deep-Dream) or independent of (Fixed) the model’s input (see Table 1) (Nguyen et al., 2016; Suzuki et al. 2017). We used the Fixed error function to simulate the high degree of spontaneity associated with complex neurological and CBS VHs, and the Deep-Dream error function to simulate the apparent dependency between visual input and hallucinatory content (i.e., the lower spontaneity) reported in psychedelic VHs (Figure 2.2).

Finally, based on our previous findings (Suzuki et al. 2017) we simulated differences in the complexity of hallucinatory content by selecting the layer within the DCNN that AM terminates. To simulate complex VHs we used the default parameters of the model, which utilises the topmost layer of the DCNN (‘fc8’) (Table 1). To simulate simple VHs we restricted information flow during AM to an arbitrary lower layer (‘Conv4’) of the DCNN, resulting in an overemphasis of the visual features extracted by these layers during training (Figure 2.3).

See supplemental material for details of the input images, minor optimisations and the number of iterations used for all simulations.

Table 1 provides a summary of the phenomenological characteristics of non-hallucinatory and hallucinatory experiences that were simulated, along with the corresponding manipulations used to simulate these characteristics.

#### 2.1.5 Image Similarity Analysis

To provide an objective method of measuring the realism of our simulations of hallucinatory phenomenology we used a commonly applied method for assessing the realism of synthetic images produced by generative models: the Inception Score (IS)(Salimans et al., 2016). To calculate the IS an image is entered into an “inception network”, a class of DCNN (e.g., GoogleNet), trained for image classification (Szegedy et al., 2014). An image produces the highest IS when it activates only a single neuron in the categorical layer of the DCNN, meaning that the features of the image converge into a single categorical label. We compared the IS between input images, benchmark, and all simulations of synthetic VHs.

To calculate the IS we used the same 5 images as in all simulations, in addition, we selected 27 random image categories from CaffeNet and used the same approach outlined in section 2.1.5 to select 27 random images (see Figure S1 for the images used). This led to a total of 32 input images. For each image we then generated synthetic examples of non-hallucinatory and hallucinatory experience using the model parameters described in section 2.1.4., resulting in six main categories: 1. Non-hallucinatory (veridical) perceptual phenomenology 2. Simple CBS VHs, 3. Simple psychedelic VHs, 4. Complex neurological VHs, 5. Complex neurological VHs, 6. Complex psychedelic VHs (see Supplemental Material for all synthetic non-hallucinatory and hallucinatory images).

To provide a baseline IS for our model we first calculated the IS for the 32 input (unaltered) images. We then calculated the IS for each of the six categories. This resulted in a single IS score between 0 and 32 for the baseline and six categories, with higher numbers denoting greater realism of the images.

### 2.2 Experimental studies: Verifying the validity of synthetic VHs

To assess the ability of our model to accurately simulate etiologically distinct VHs, we conducted two separate studies: (i) an online survey that targeted participants with recent experience of classical hallucinogens, and (ii) a semi-structured series of phenomenological interviews in which patients with Lewy Body Dementia, Parkinson’s Disease or Charles Bonnet Syndrome described their VHs in detail.

#### 2.2.1 Online Psychedelic Survey

This experiment was designed to assess how closely participants’ chosen psychedelic experiences matched the synthetic VHs produced by our model. We reasoned that if our simulations were successful in producing representative examples of psychedelic experience, participants would be more likely to select synthetic psychedelic VHs, compared to neurological VHs, as most closely matching their experience. The experiment was carried out in accordance with approved guidelines provided by the University of Sussex, Research Ethics Committee, and was pre-registered using OSF (https://osf.io/qu8xe). Here, we report abbreviated methods, for a full description of the experiment see supplemental methods.

##### Participants

Eighty-one participants completed the online survey who reported having taken a classical hallucinogen (LSD, psilocybin, or DMT) within the previous 12 months.

##### Procedure

The experiment consisted of collection of background information, an image selection task, and assessment of hallucinatory phenomenology. It began by asking eligible participants to select a particular psychedelic experience from within the last 12 months, that they would use to answer all the questions in the survey, the type of classical hallucinogen this experience related to and the subjective potency of this experience using a scale from 0 (not potent at all) to 5 (as potent as my most intense psychedelic experience).

Participants then completed 5 practice trials of the image selection task. Each trial of the image selection task presented 6 randomly selected synthetic VHs images (3 neurological and 3 psychedelic) participants were asked to select which image (if any) that most closely resembled their chosen psychedelic experience (https://osf.io/nr4ke/files/osfstorage) (see supplemental methods for more detail).

The main experiment used the same trial structure as the practice session but consisted of a total of four blocks, each with 32 trials. Two blocks contained images simulating simple VHs (3 CBS and 3 psychedelic), and two blocks contained images simulating complex VHs (3 neurological and 3 psychedelic). Within each block, 2 catch trials were placed in random positions that contained task irrelevant instruction (e.g., click the top-right image), to maintain participants’ attention.

Following the main experiment, participants were asked to indicate if their chosen psychedelic experience contained simple, complex or both types of hallucinatory content. They were then asked separately about the veridicality (from 1 (identical) to 10 (completely different)), and the spontaneity (scale from 1 (completely dependent) to 5 (completely independent)) of each type of hallucinatory content in relation to their existing visual experience.

##### Pre-registered Analysis

We followed the analysis plan and exclusion criteria described in the pre-registration document (https://osf.io/qu8xe). We excluded participants who skipped more than 75% of the trials, those who had an average reaction time of less than one second, and those who did not meet the 1-minute time limit for more than 50% of all trials. In addition to the pre-registered exclusion criteria, we set post-hoc exclusion criteria in which participants who answered four or more of the eight catch trials incorrectly were excluded.

To investigate if synthetic psychedelic VHs were phenomenologically similar to psychedelic experiences, we used a one-tailed, one-sample t-test using the test variable of the sample mean and a chance level of 0.5 (50%) to compare the ratio of responses between psychedelic and neurological synthetic VHs for both simple and complex blocks separately. A resulting p-value equal to or less than 0.05 (P ≤ 0.05) was interpreted as evidence that participants selected psychedelic synthetic VHs at a higher-than-chance frequency. This standard t-test was accompanied with a Bayesian t-test. A Bayes Factor (BF) greater than 3 was taken to indicate sensitive evidence for the hypothesis that participants selected simulations of psychedelic VHs at a higher-than-chance frequency, while a BF<1/3 was taken to indicate sensitive evidence for the null hypothesis. BFs between these two thresholds (>1/3 but <3) were taken as evidence that the data was insensitive (Dienes, 2014; Jeffreys, 1939).

##### Exploratory Analysis

We performed two exploratory analyses which were not included in the pre-registration.

First, to explore if any of the other image generation parameters used to produce the synthetic VHs affected how representative of psychedelic experience they were, we conducted two separate ANOVAs (simple VHs and complex VHs) for the most frequently selected hallucination type (either psychedelic or neurological VHs) based on the pre-registered results, using image selection frequency as the dependent variable. For complex VHs we conducted a 3 x 2 ANOVA of image selection frequency, with factors iteration number (10, 100, or 1000) and error function (WTA or Fixed). For simple VHs we conducted another separate 3 x 2 ANOVA of image selection frequency, with factors iteration number (10, 100, or 1000) and target layers (conv3 or conv4).

Second, we examined the correlation across participants between the reported potency of the chosen psychedelic experience and the number of iterations used to generate the synthetic VHs again for the most selected hallucination type (either psychedelic or neurological), reasoning that those participants who had reported a higher potency psychedelic experience would be more likely to select synthetic VHs produced by a higher number of iterations. First, for each participant, we ran a linear regression across all stimuli, plotting the iteration level of each stimulus against how frequently that stimulus was selected. The slope value (coefficient) of each regression reflects the iteration level preference for a participant, with a positive slope indicating a tendency to select a higher iteration value. We then ran a one-tailed bivariate correlation (Pearson’s coefficient) across participants using the variables of ‘iteration preference’, taken from the slope of the regression, and potency rating (0-5).

#### 2.2.2 Clinical semi-structured phenomenological interview

Twenty-two participants (13 female) took part in a semi-structured interview designed to enquire about the visual phenomenology associated with CBS and neurological VHs (PD and LBD, see S5)(mean age = 67 years SD 15.9). Nine participants had received a formal diagnosis of Parkinson’s Disease (PD), three a formal diagnosis of Lewy Body Dementia (LBD), and ten reported visual loss resulting in a formal diagnosis of Charles Bonnet Syndrome (CBS). This experiment was carried out in accordance with approved guidelines provided by the University of Sussex, Research Ethics Committee.

All participants were interviewed by author D.J.S by video conference call (zoom etc.) or telephone, and each interview lasted no longer than 45 minutes. After an introduction, and verbal consent was provided, the interview began by collecting background information about general features of participants’ VHs, such as how long they had experienced visual hallucinations, how frequently they occurred, how long on average each VH lasted, the differing types of content of their VHs, the complexity of their VHs (simple/complex) and if they had ever confused their VHs as being real (reality monitoring).

Each participant was then asked to describe in as much detail as possible their most recent (target) experience of a VH. We used open-ended questioning to allow participants to answer in detail. For each target VH, the spontaneity of the VH was assessed by the context of the description. A VH was interpreted as being spontaneous if it was described as being a transformation of pre-existing aspects of a visual scene, as compared to occurring in the absence of any corresponding pre-existing content. If there was any ambiguity in participants’ descriptions regarding spontaneity, the interviewer guided the participant towards reporting on this aspect of their VH, by asking them specifically if their VHs occurred in the absence of pre-existing content within the visual scene.

Participants were then asked to rate the veridicality of the VH: “On a scale of 1-10, 10 being identical to normal visual experiences, 1 being not at all like normal visual experience, how would you rate the visual quality of this VH? “. For CBS participants in which no vision was preserved, they were asked to compare the veridicality of the target VH to memories of their visual experience before the onset of eye disease. This procedure was repeated with other instances of VHs that the participant reported, up to a maximum of 5 descriptions.

Following this portion of the interview, each participant was shown a 5 x 10 grid of images (see Figure 3) displaying examples of appropriate synthetic VHs. The first column displayed the 5 unaltered input images used in all simulations. Each successive column displayed synthetic complex neurological or CBS VHs (dependent on the participant’s condition) following 5, 10, 50, 100, 200, 400, 600, 800 and 1000 iterations. The participants were instructed: ‘please select the column, if any, that displays the closest visual similarity to your experience of visual hallucinations’.

**Figure 3.**
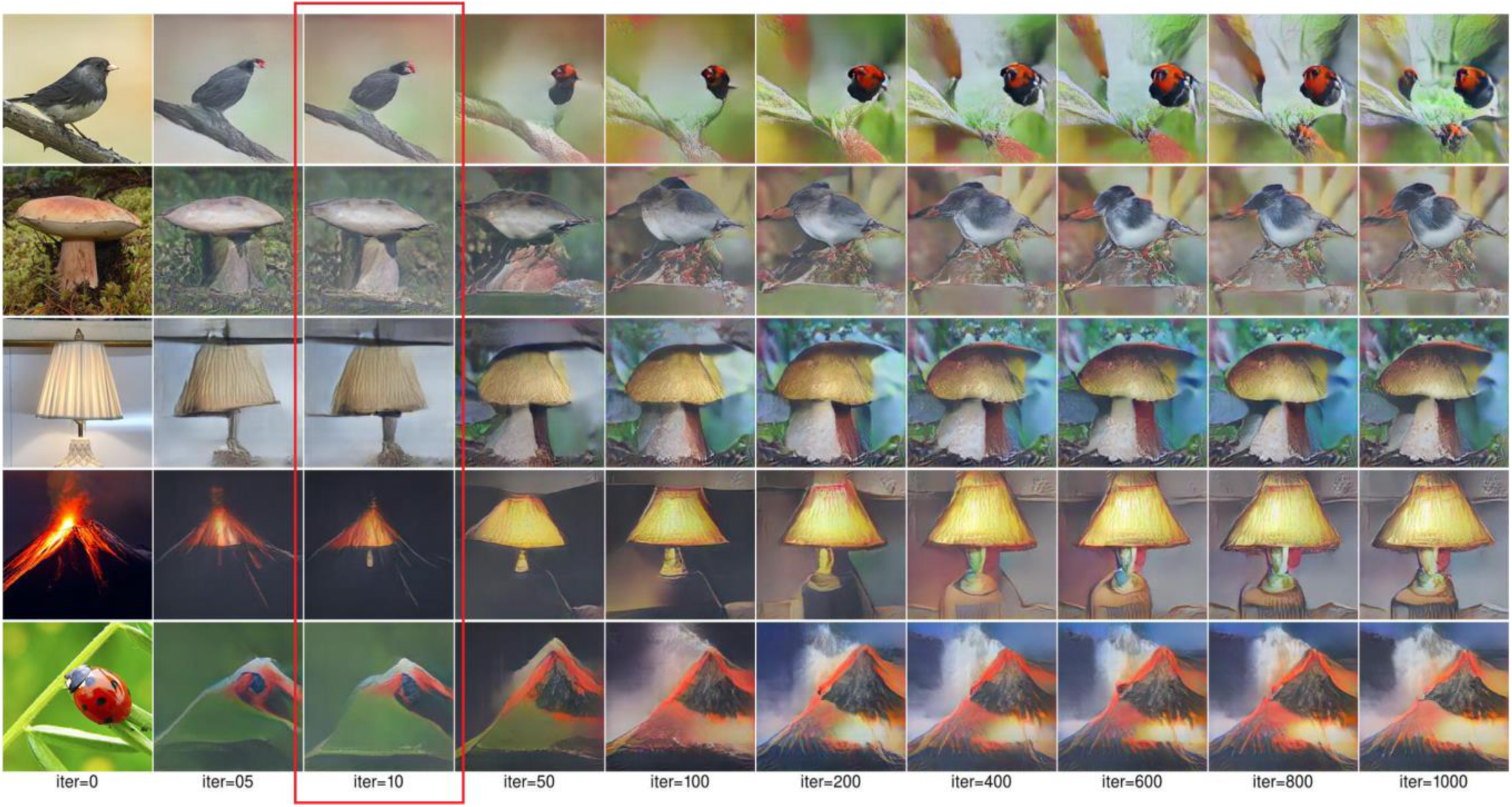
Synthetic complex neurological VHs used to assess if they were representative of neurological and CBS hallucinatory phenomenology during the phenomenological interview. Participants were asked to choose which column, if any, was similar in visual quality to their experience of complex VHs. The red box highlights the nearest whole iteration number (average value = 10.8) that neurological and CBS participants rated on average as being closest in visual quality to their complex VHs. Parameters used to generate these synthetic VHs are identical to those used in Figure 5.

If a participant reported simple VHs, they were also shown a similar grid of simple synthetic CBS VHs (see Figure 4). Three CBS participants were too visually impaired to perform this task.

**Figure 4.**
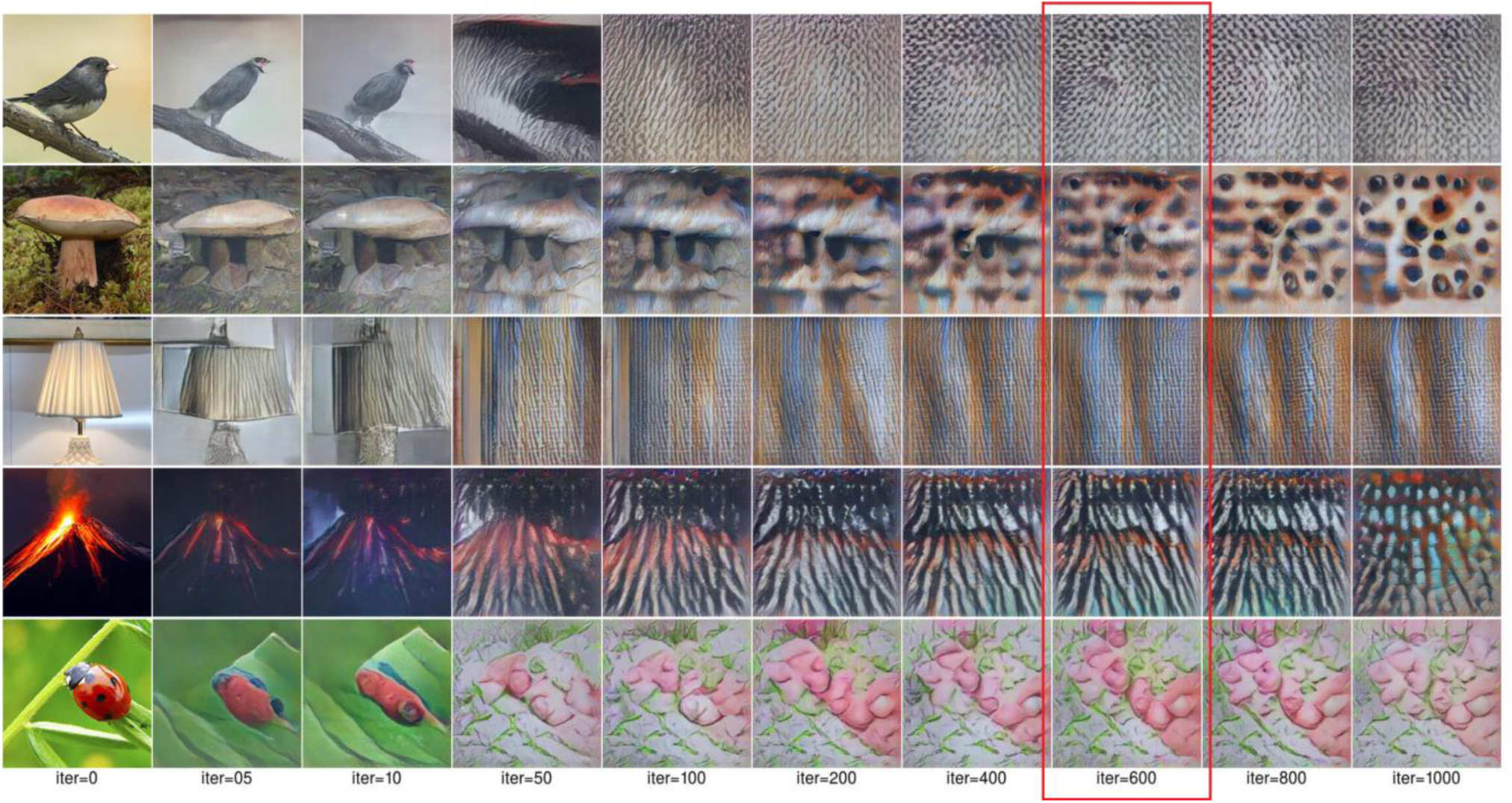
Synthetic simple CBS VHs used to assess if they were representative of CBS hallucinatory phenomenology in the phenomenological interview. Participants were asked to choose which column, if any, was similar in visual quality to their experience of simple VHs. The red box highlights the nearest iteration number (average value = 650) that CBS participants rated on average as best characterising the visual quality to their simple VHs. The parameters used to generate these synthetic VHs are identical to those used in Figure 6.

## 3 Results

We simulated three distinct aspects of clinical and psychedelic visual hallucinatory phenomenology – their complexity, veridicality and spontaneity – by manipulating a pre-trained coupled DGN-DCNN model. These distinct aspects were selected to reflect the reported phenomenology of VHs experienced by people with certain neurological conditions, by people with visual loss, and by neurotypical people following the ingestion of classical psychedelics (see Table 1). We assessed the output of the model objectively by comparing the Inception Scores from our benchmark simulation of non-hallucinatory experience to all other simulations of hallucinatory phenomenology. The Inception Scores of all the simulation results are presented in Table 2. We also conducted both questionnaire and semi-structured interview surveys with people from each of the above three groups to assess the subjective match between model output and reported experience.

**Table 2.** Comparison of Inception Scores for 32 arbitrary input images and synthetic outputs of the model for each simulation. A higher number denotes a greater degree of visual realism of the image set. Note that the maximum Inception Score is derived from the total number of images used in the calculation i.e., 32.

| Image Type | Inception Score<br>(IS) |
| --- | --- |
| Input images | 28.54 |
| Non-Hallucinatory Perception<br>(Benchmark) | 22.59 |
| Neurological | 19.92 |
| CBS Complex | 22.40 |
| CBS Simple | 6.68 |
| Psychedelic Complex | 5.52 |
| Psychedelic Simple | 3.42 |

### 3.1 Simulating non-hallucinatory perceptual phenomenology

To provide a benchmark of model performance, we first simulated non-hallucinatory (veridical) perceptual phenomenology. Note that if the performance of the model was perfect in simulating non-hallucinatory perceptual phenomenology, then the input and output should be identical. Figure 5 shows the procedure and representative results from the benchmark simulation.

**Figure 5.**
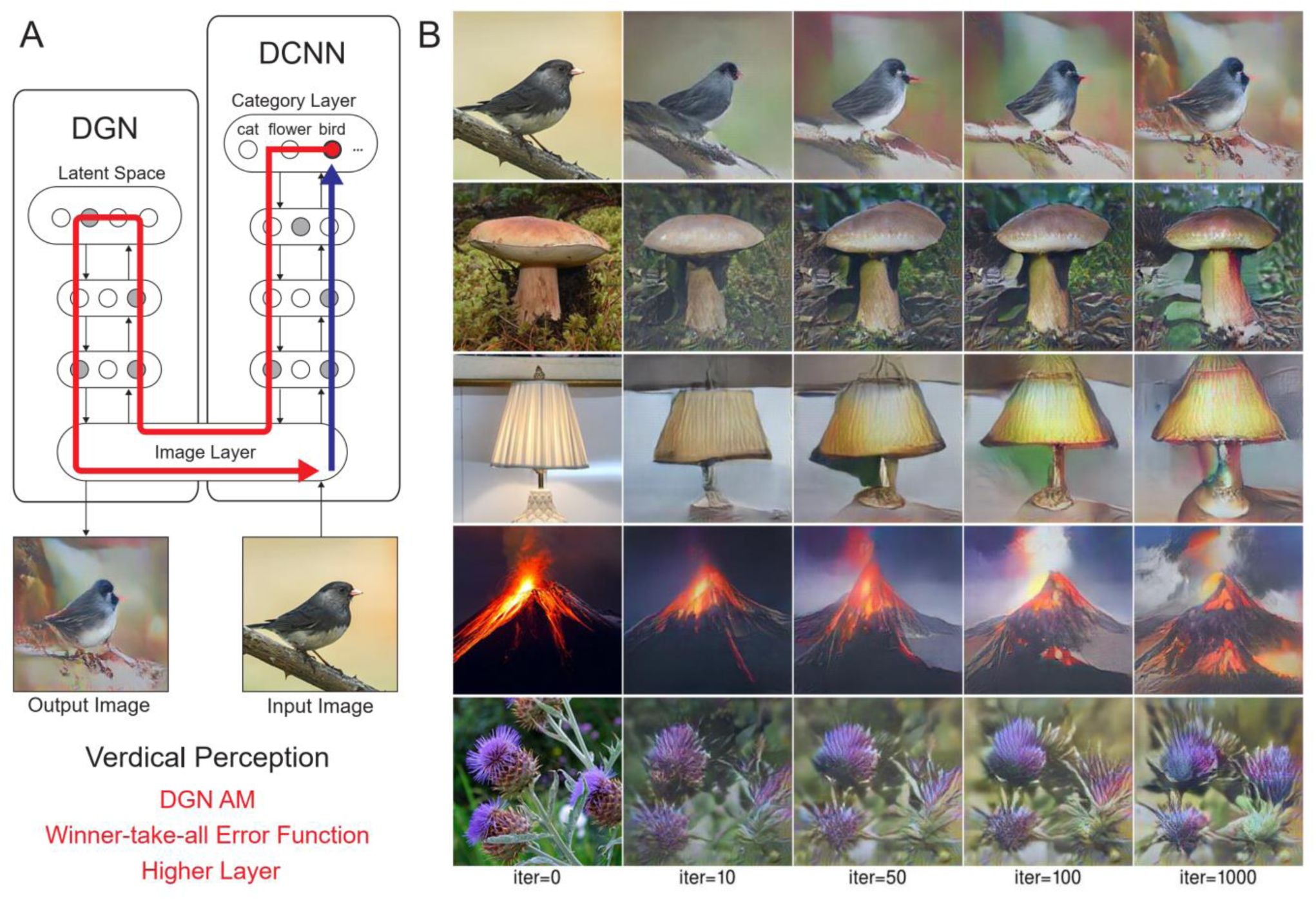
Model architecture and outputs simulating benchmark (non-hallucinatory) perceptual phenomenology. A. Schematic of model architecture and information flow for a single iteration. An initial arbitrary input image is passed forward through the DCNN, which extracts the visual features of the image across the layers of the network (Blue arrow). Using the winner-take-all error function, the neuron that responds maximally to the input image neuron is selected within the highest categorical layer of the DCNN. The categorical information held within the selected target neuron is passed to the DGN, which updates the latent space of the DGN and creates a new synthetic image that maximises the activity of the selected target neuron (Red arrow). B. The initial input images (5 images, left column) and visualisations of the network following 10, 50, 100 and 1000 iterations.

Using the Inception Score to measure the realism of the benchmark simulation we found that the synthetic outputs of this simulation displayed the highest Inception Score (22.59) of all the reported simulations when compared to the (unaltered) input images (28.54). We used these visualisations and Inception Score as a benchmark with which to assess the veridicality of further simulations.

### 3.2 Simulating neurological visual hallucinations

Next, we simulated the perceptual phenomenology associated with complex neurological VHs (PD and LBD).

Simulating complex neurological VHs we found the model’s visualisations displayed high veridicality (Figure 6), producing a relatively high Inception Score (19.92) when compared to the benchmark (22.59). In addition, due to the use of the Fixed error function and selection of an experimenter-determined target neuron, the resulting visualisations were not at all related to the input image. Together, these synthetic VHs display the key phenomenological characteristics of high veridicality, spontaneity and complexity that are typical of complex neurological VHs.

**Figure 6.**
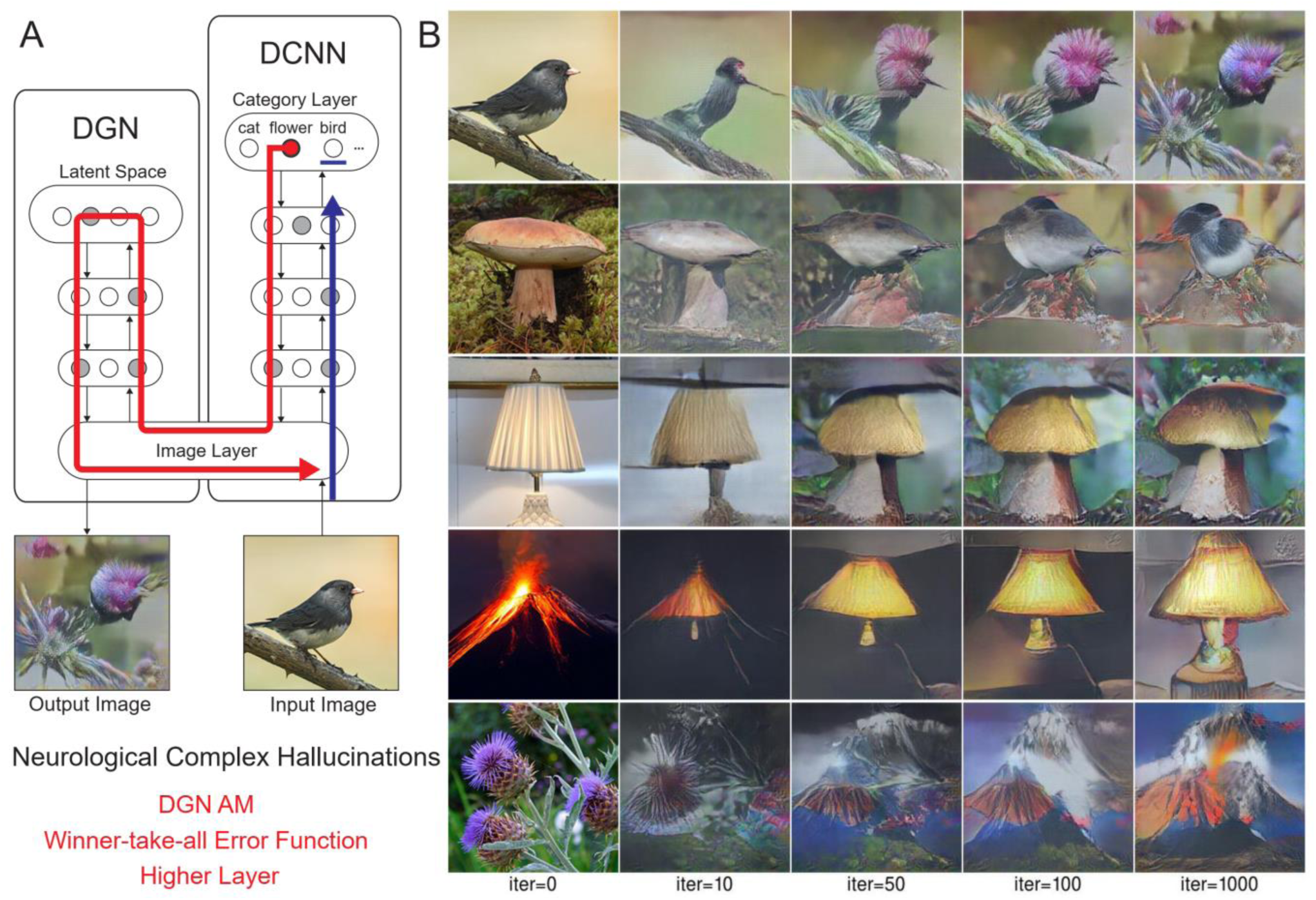
Model architecture and synthetic VHs simulating neurological complex VHs. A. Schematic of model architecture for a single iteration. An arbitrary input image (e.g., bird), is passed forward through the DCNN (Blue arrow). Irrespective of the input image, an experimenter-determined target neuron is selected within the DCNN (e.g., flower) using the Fixed error function. This information is passed to the DGN and updates the latent space so that the generated image increases the activation of the target neuron (Red arrow). B. The initial input images (left column) and synthetic VHs of the network following 10, 50, 100 and 1000 iterations. From top to bottom, the experimenter-determined target neuron was Flower, Bird, Mushroom, Lamp and Volcano.

### 3.3 Simulating visual hallucinations due to visual loss

Next, we simulated the perceptual phenomenology of simple and complex CBS VHs by introducing a central ‘blur’ to the input image (see Methods and Figure 7) and keeping all other parameters the same as complex neurological VHs (Figure 6). We found that the resulting synthetic VHs displayed high veridicality, which was confirmed by a high Inception Score (22.40) when compared to both our benchmark (22.59) and neurological complex VHs (19.92). Together, these synthetic VHs are in line with typical reports of complex VHs in CBS (Abbott et al. 2007; ffytche and Howard 1999; Menon et al. 2003; Schultz et al. 1996; Teunisse et al. 1996), displaying both high veridicality, spontaneous content and complexity.

**Figure 7.**
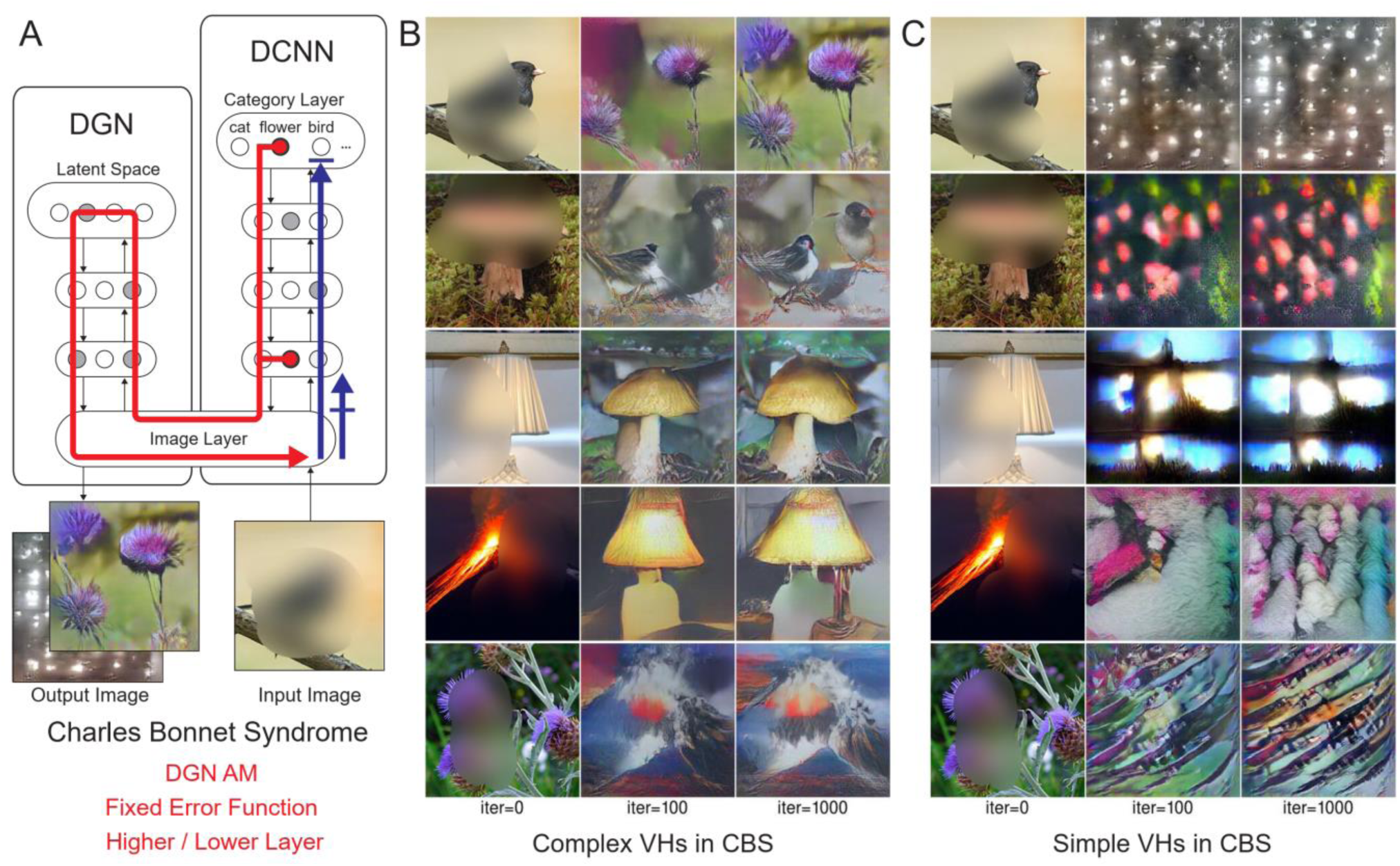
Model architecture and synthetic VHs simulating simple and complex VHs CBS. A. Schematic of the model architecture used in a single iteration. An arbitrary input image (e.g., bird) with representative features of the visual deficits associated with CBS (central blur), is passed forward through the DCNN (Blue arrow). Irrespective of the input image an experimenter-determined target neuron is selected in the DCNN output layer (e.g., flower) using the Fixed error function. With each iteration the DGN generates a new image that maximally activates the target neuron (Red arrow). For simple CBS VHs, we restricted the level within the DCNN that AM terminates to a lower layer (Conv4). B. Simulations of complex CBS VHs, iterating from an input image (left column) and outputs following 100 and 1000 iterations. From top to bottom, the Fixed error function maximises the activity of the DCNN target neuron representing flower, bird, mushroom, lamp and volcano. C. Simulations of simple CBS VHs iterating from an input image (left column) and outputs following 100 and 1000 iterations. Irrespective of the input image an experimenter-determined target neuron is selected in the lower layer (Conv4) of the DCNN using the Fixed error function. With each iteration, the DGN alters the image based on the learnt features contained within the selected target neuron in the Conv4 layer of the DCNN.

To simulate simple CBS VH, we restricted the level within the DCNN that AM terminates to a lower layer (conv4) and allowed the DGN to synthesise new images based on the activity of a randomly selected target neuron within this layer (Figure 2.3). The resulting synthetic VHs contained low-level colours and textures associated with natural real-world objects, similar to the flashes of light, abstract shapes and repeating patterns commonly reported in simple CBS VHs (Abbott et al. 2007; ffytche and Howard 1999; Menon et al. 2003; Teunisse et al. 1996). Due to the predominance of low-level visual content of these visualisations the resulting IS was as expected lower (6.68) than both benchmark and other simulations of complex VHs.

### 3.4 Simulating psychedelic visual hallucinations

Using the same model parameters as in our previous simulations of psychedelic hallucinatory phenomenology (Suzuki et al., 2017), we were able to simulate the reduced veridicality and dependency on sensory input (reduced spontaneity) associated with complex psychedelic VHs. The use of a consistent set of input images across all simulations in this paper enables the comparison of synthetic psychedelic VHs with aetiologically distinct synthetic VHs.

Compared to benchmark, neurological and CBS synthetic VHs, simulations of complex psychedelic VHs displayed low veridicality, reflected by a low Inception Score (5.52). In addition, the complex hallucinatory content within these synthetic VHs can be seen as being driven by visual ‘seeds’ within the input image (Figure 8). These synthetic VHs also display hallucinatory transformations of the input image, for example, an input image of a mushroom or flower (2nd and 5th rows) is transformed into ‘fish’ or ‘bird-like’ hallucinatory content that still somewhat conforms to the global structural properties of the input image (Figure 8). Together, these synthetic VHs are in line with anecdotal reports of psychedelic complex VHs displaying lower veridicality and being driven by visual ‘seeds’ within an observed scene.

**Figure 8.**
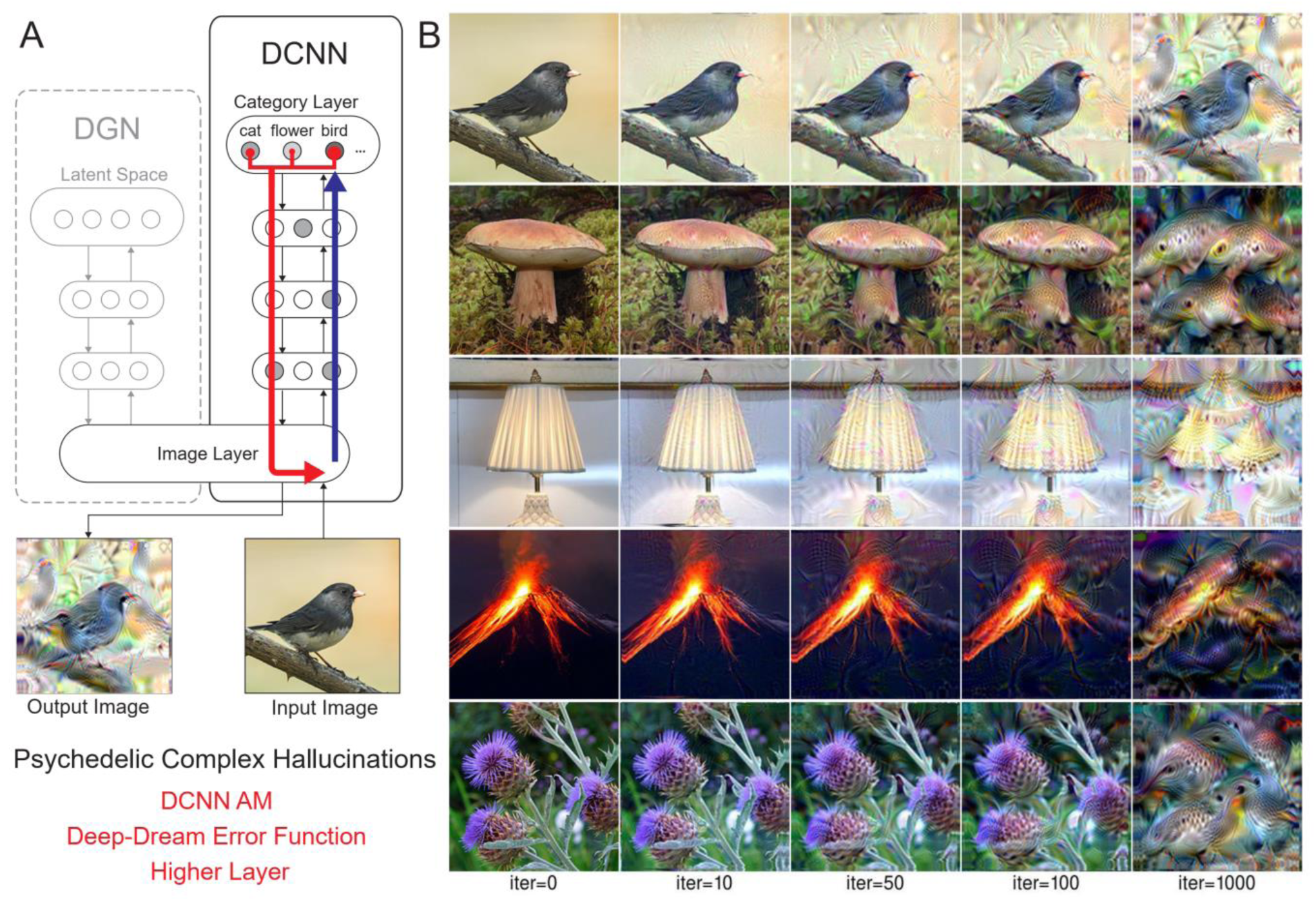
Model architecture and synthetic VHs simulating complex psychedelic VHs. A. Schematic of the model architecture used in a single iteration, using only the DCNN and Deep-Dream error function. An initial input image is passed forward through the DCNN extracting the visual features of the image across the layers of the network. Using the Deep-Dream error function, errors are returned for all neurons within the highest layer of the DCNN that were activated by the input image. These errors are transmitted via backpropagation to alter the colour of each pixel within an input image to maximise activity within the activated target neurons. B. Simulations of complex psychedelic VHs. The initial input images (left column) and synthetic VHs of the network following 10, 50, 100 and 1000 iterations. Note that this architecture is identical to that used in Suzuki et al., (2017).

Finally, to simulate simple psychedelic VHs, we used the same parameters as above but restricted the level within the DCNN that AM terminates (Conv4) (see Figure 9). In contrast to simulations of simple CBS VHs (Figure 8), the resulting visualisations prominently included geometric shapes, patterns and rhythmic kaleidoscopic imagery similar to those typically reported during psychedelic experiences (Bressloff et al., 2001; Cowan, 2015; Díaz, 2010; Nichols, 2016; Tyler, 1978). The marked ‘hallucinatory’ quality of these visualisations was reflected by a reduced Inception Score (3.42), as compared to simple CBS VHs (6.68). Similar to our simulations of complex psychedelic VHs, these visualisations were also transformations of existing sensory input. As shown in Figure 8, some of the geometric features within each visualisation are clearly driven by the corresponding properties of the input image.

**Figure 9.**
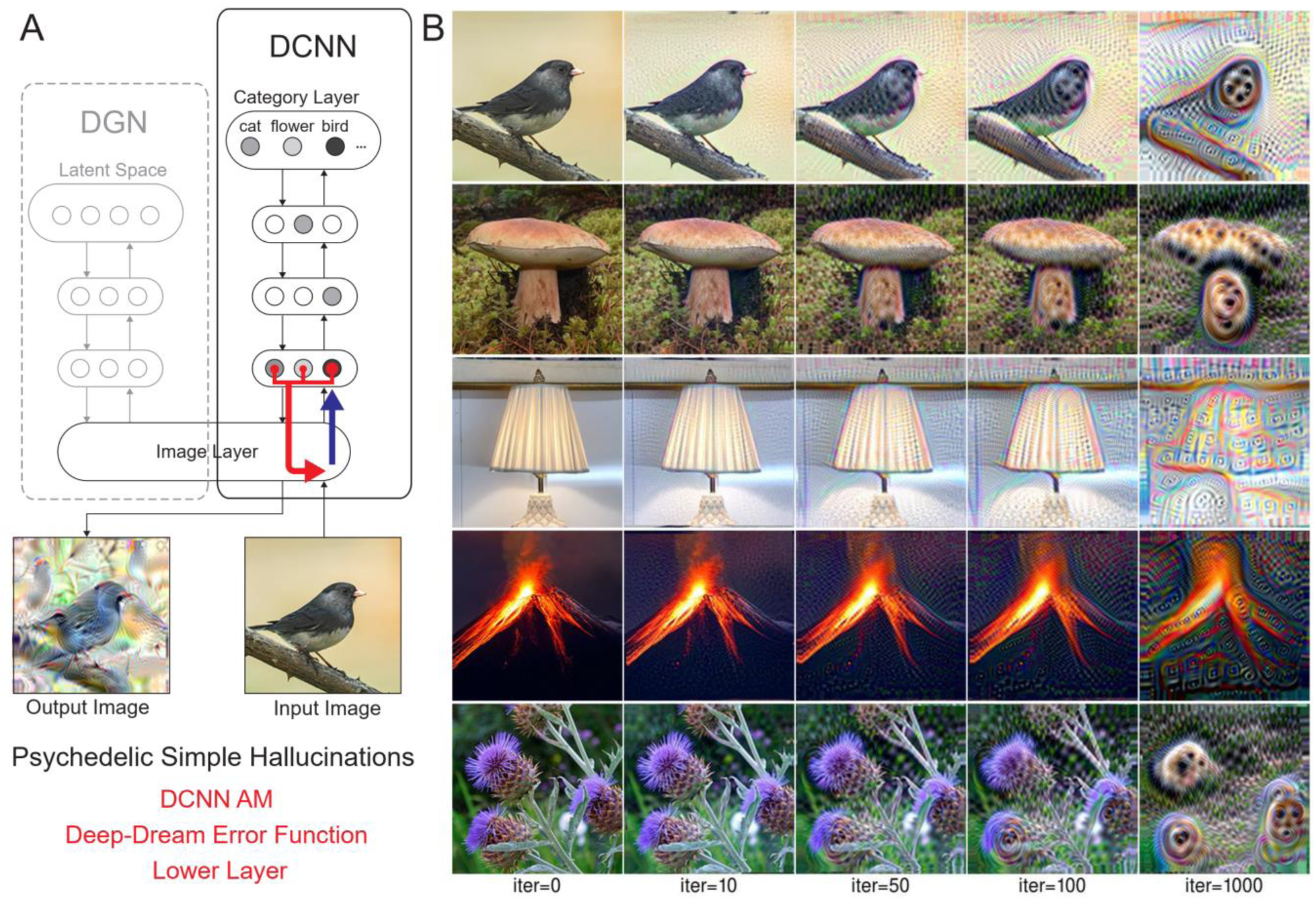
Model architecture and synthetic VHs simulating simple psychedelic VHs. A. Schematic of the model architecture for a single iteration: the level that DCNN-AM terminates was restricted to a lower layer (Conv4). An initial input image is passed forward through the DCNN. Using the Deep-Dream error function, errors are returned for all neurons within the Conv4 layer of the DCNN that were activated by the input image. These errors are transmitted via backpropagation to alter the colour of each pixel within an input image to maximise activity within the activated target neurons. B. Input images (left column) and visualisations of the network restricted to the Conv4 layer of the DCNN, following 10, 50, 100 and 1000 iterations. Note that this architecture is identical to that used in Suzuki et al., (2017).

### 3.5 Clinical semi-structured phenomenological interview

A semi-structured phenomenological interview was used to enquire about the visual phenomenology associated with CBS and neurological VHs (see Table 3 and https://osf.io/nr4ke/files/osfstorage). Assessing the veridicality of complex VHs, we found that 92% of neurological (PD and LBD) and 90% of CBS patients reported that their VHs appeared ‘as real’ or ‘similar to’ their normal visual experiences. In terms of spontaneity, 18 out of 20 participants (2 CBS participants were severely visually impaired) reported that their VHs always occurred spontaneously, in the absence of sensory cues, with two PD participants reporting that infrequently their VHs could be described as transformations of existing visual information within the observed scene. For example, one PD participant described an occasion in which their slippers had turned into rats that ran across the bathroom floor. Examining the complexity of the reported hallucinatory content, we found that 92% of PD and LBD participants reported that their VHs were complex in nature, with only one participant reporting both simple and complex VHs. In contrast, 90% of CBS participants reported experiencing both simple and complex VHs, with only one participant reporting only complex VHs.

**Table 3.** Results of semi-structured clinical interviews, showing demographic information and characteristics of PD, LBD and CBS VHs.

| Demographics | PD/LBD | CBS | Frequency of VHs | PD/LBD | CBS |
| --- | --- | --- | --- | --- | --- |
| N = | 12 | 10 | Daily | 58% | 60% |
| Age | 69±8.61 | 72.1±21.1 | Weekly | 17% | 0% |
| Gender | M=7 / F=5 | M=2 / F=8 | Monthly | 25% | 40% |
| Veridicality |  |  | How long VHs have occurred |  |  |
| Normal visual experience (9-10) | 67% | 30% | <1 year | 42% | 0% |
| Similar (5-8) | 25% | 60% | 1-10 years | 50% | 70% |
| Less Clear (<5) | 8% | 10% | >10 years | 8% | 30% |
| Spontaneity |  |  | Duration of VHs |  |  |
| Spontaneous | 83% | 100% | 1-5 s | 8% | 0% |
| Visual transformation | 17% | 0% | 5-60 s | 33% | 30% |
|  |  |  | 1-60 min | 58% | 70% |
| Complexity of VHs |  |  | Content |  |  |
| Simple only | 0% | 0% | Adults | 92% | 90% |
| Complex only | 83% | 10% | Children | 33% | 10% |
| Both | 17% | 90% | Animals | 50% | 50% |
|  |  |  | Scenes | 8% | 10% |
| **Reality Monitoring** |  |  | Other objects | 0% | 30% |
| Preserved | 50% | 75% |  |  |  |
| Not Preserved | 25% | 25% |  |  |  |
| Variable | 25% | 0% |  |  |  |

Next, to assess our model’s ability to produce representative visualisations of neurological and CBS VHs we showed participants a range of synthetic neurological or CBS VHs generated using an increasing number of iterations and asked them to select which (if any) was the most representative of their hallucinatory experience (Figure 3). Due to severe visual or cognitive impairment three CBS and two LBD patients were unable to complete this task. Supporting the validity of our synthetic VHs, we found that all participants were able to find a synthetic VH that they reported as being representative of their hallucinatory experience. Next, we predicted that due to the high reported veridicality of neurological and CBS VHs that participants would select synthetic VHs with a low number of iterations, i.e., those that were visually similar to the input image. Pooling the images selected by PD and LBD participants and averaging each image’s iteration number resulted in an average iteration value of 3 (N = 10; SD = 4.3), suggesting that for this group their VHs displayed high veridicality. In contrast, we found an average iteration value of 18.6 for CBS complex VHs (N = 7, SD = 36.1), suggesting that for these participants their complex hallucinatory experience did not display as high a degree of veridicality as neurological patients. Pooling data across both groups resulted in the synthetic VHs produced following 10 iterations as being the closest representation of both neurological and CBS complex VHs (Figure 3).

In contrast to complex VHs, we predicted that participants who experienced simple VHs would select simple synthetic VHs with a high number of iterations, due to the abstract low-level visual features requiring more iterations to develop in the synthetic VHs. Indeed, on average, the single PD and five CBS patients who experienced simple VHs selected synthetic VHs following 650 iterations (N = 6; SD = 333.1) as being the most accurate representation of their simple VHs (Figure 4).

### 3.6 Psychedelic survey

#### 3.6.1 Pre-registered analyses

The number of participants for this study did not reach the pre-registered sample size of 200, therefore the following analysis is exploratory. No participants were excluded based on the pre-registered exclusion criteria for this study. Applying the post-hoc exclusion criteria (less than 4 correct answers to the catch trials), four participants were excluded (4.9% of all participants) out of the eighty-one participants who completed the online survey.

Out of the remaining seventy-seven participants, 46 indicated that their reports related to the ingestion of psilocybin, 24 LSD, and 7 DMT. When asked to rate the subjective potency of their hallucinatory experience on a scale of 0 (not potent at all) to 5 (as potent as my most intense psychedelic experience), average potency ratings were: psilocybin 3.6 (SD=1.28), LSD 3.5 (SD=1.1) and DMT 4.3 (SD=0.76) (Figure 10).

**Figure 10.**
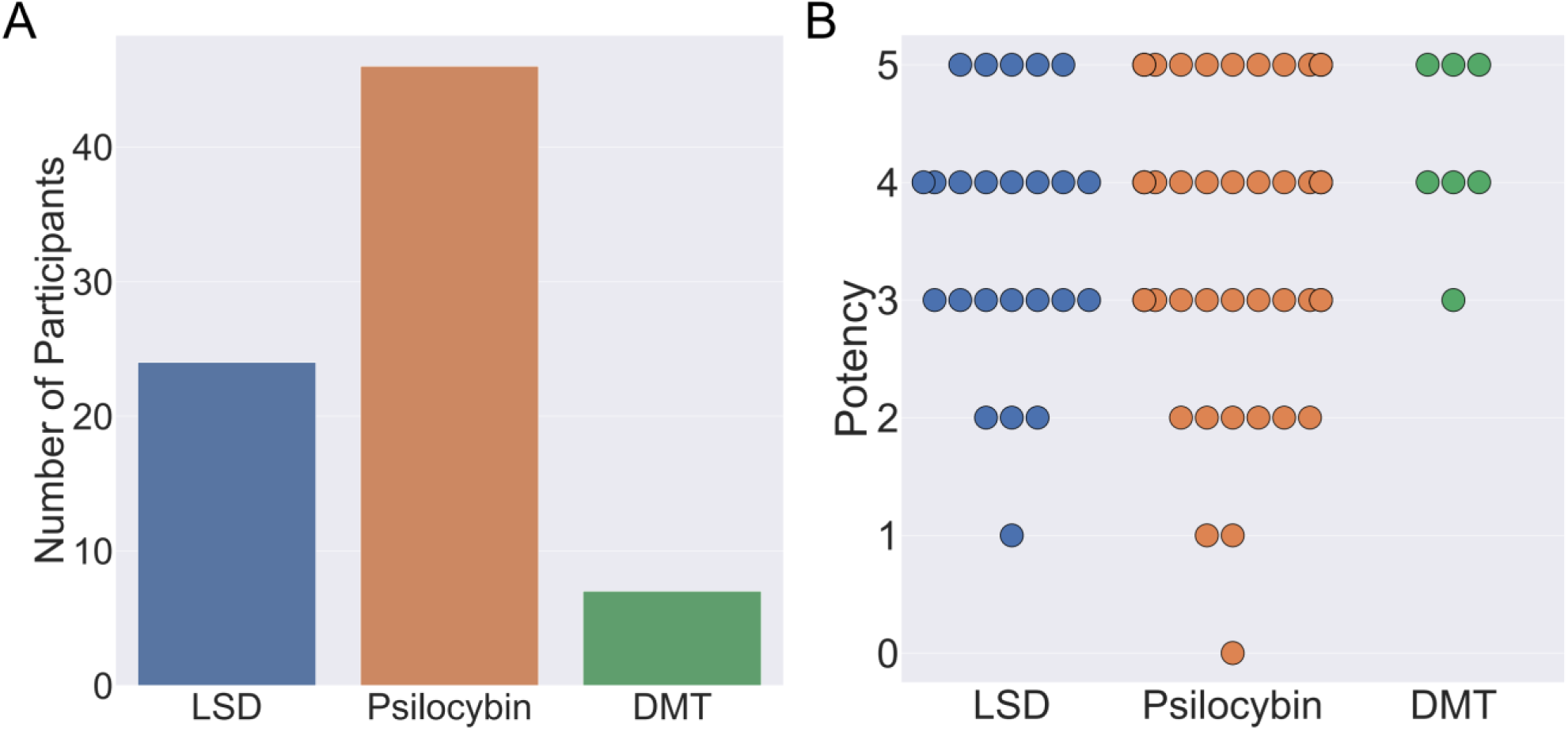
A. Classical hallucinogen taken by participants during their chosen psychedelic experience (left). B. Reported potency of participants chosen psychedelic experience on a scale of 0 (not potent at all) to 5 (as potent as my most intense psychedelic experience ever). Each circle denotes an individual participant response.

To investigate if synthetic psychedelic complex VHs were representative of psychedelic experience, we first examined the results of the image selection task from the psychedelic survey (Figure 11). We found that participants who reported complex VHs chose synthetic psychedelic VHs significantly more frequently than synthetic neurological VHs (one-tailed t-test, t(76)=13.02, p <0.01, Cohen’s d=3.80, BF10 = 1.92×1018). Similarly, participants who reported simple VHs selected synthetic psychedelic VHs significantly more frequently than synthetic neurological VHs (t(76)=12.26, p<0.01, Cohens’ d=3.75, BF10= 9.51×1016). These results suggest that, for this sample, synthetic psychedelic VHs were representative of both simple and complex psychedelic phenomenology.

**Figure 11.**
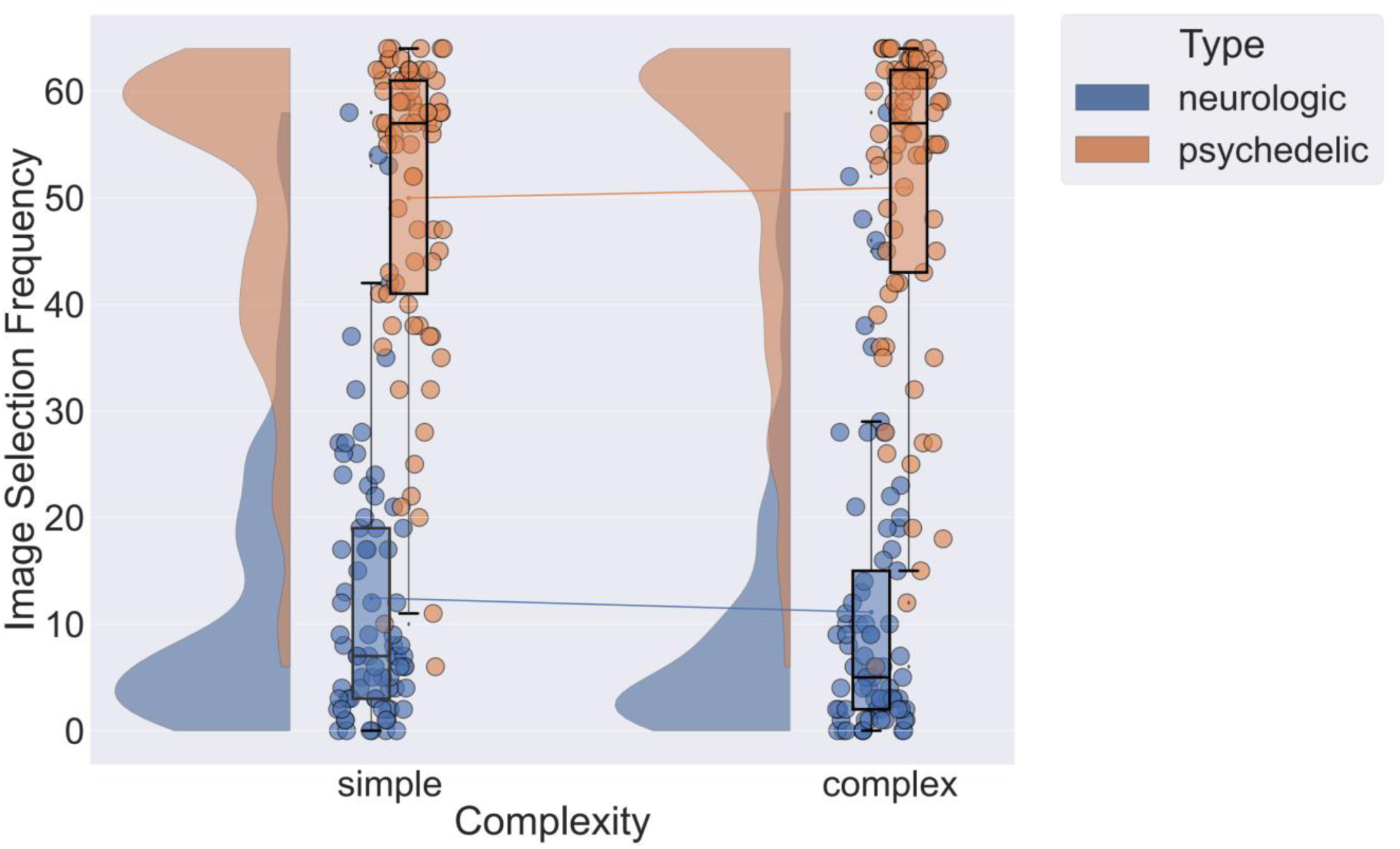
A raincloud plot showing the number of neurological or psychedelic images selected by participants in the image selection task for simple and complex blocks. For each participant responses are averaged across two simple and two complex blocks (note there were a total of 32 neurological and 32 psychedelic images per block). Each circle indicates the total numbers of images selected by a participant for each class of image. Lines between plots denote the mean frequency of response for neurological or psychedelic images averaged across participants for simple and complex blocks.

#### 3.6.2 Exploratory analyses

Next, to explore if the additional parameters used in the generation of the synthetic psychedelic VHs - the number of iterations, and DCNN level where AM terminated - affected the likelihood of them being selected as representative of participant’s psychedelic experience we compared these factors for each category of synthetic VH: complex-psychedelic, and simple-psychedelic using 2 separate ANOVAs. For complex VHs the factors used in the ANOVAs were iteration level (3) and AM Type (2). For simple VHs the factors used were iteration level (10,100,1000) and the layer of DCNN (2). Confusion matrices of average image selection frequency for each parameter are shown in Figure 12.

**Figure 12.**
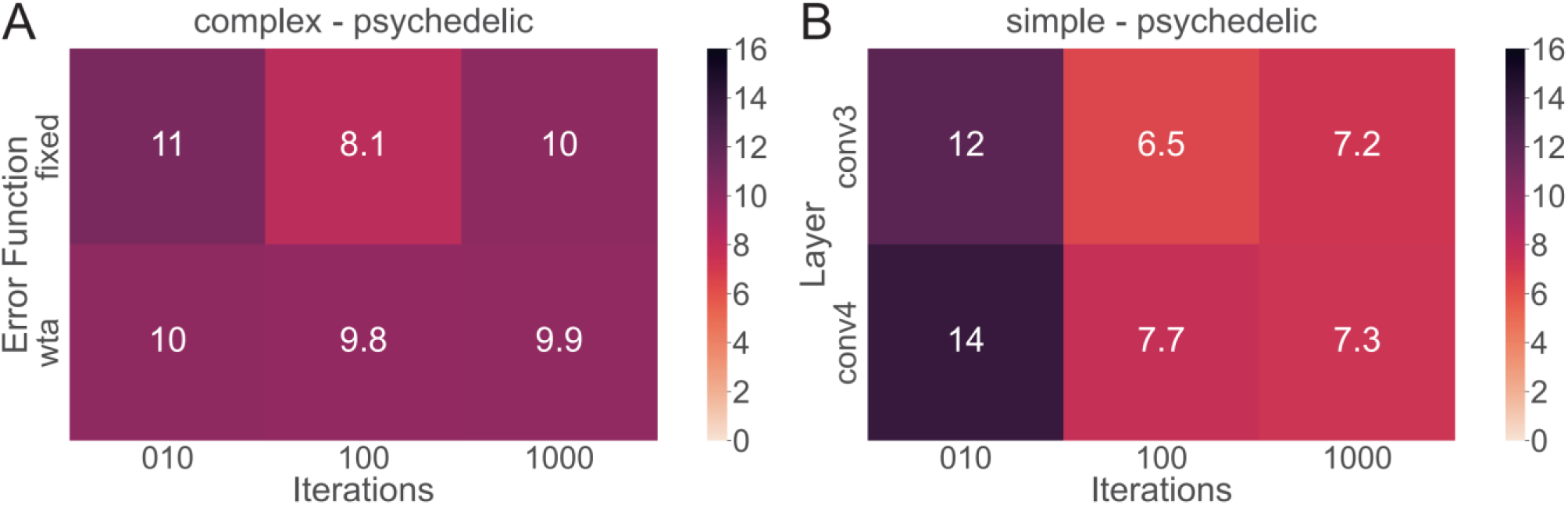
Confusion matrices showing the number of neurological or psychedelic images selected by participants for iteration number (10,100,1000), error function (Fixed, WTA) or layer at which DCNN-AM terminates (conv3, conv4). A. Confusion matrix showing the effects of error function and the number of iterations on image selection frequency for synthetic complex psychedelic VHs. B. Confusion matrix showing the effects of layer and the number of iterations on image selection frequency for synthetic simple psychedelic VHs. All matrices display the selection frequency averaged across participants and the two respective blocks (simple, complex) for each class of image. Note that the maximum number of each cell is 16.

Results of the ANOVAs revealed a significant main effect for iteration level only for Simple-psychedelic VHs (F(81,2)=29.28 p<0.01, η2 =0.12) (see Supplemental Material for all ANOVA results). Additional post-hoc tests revealed that the ratio of responses was significantly higher for 10 compared to both 100 (t(76)=6.82, pbonf<0.01, Cohen’s d=0.77) and 1000 iterations (t(76,2)=6.41, pbonf<0.01, Cohen’s d=0.70) (see Figure 12), suggesting that for these participants the majority of their simple VHs were relatively subtle in nature, consisting of low-level visual distortion and not the colourful kaleidoscopic patterning typically reported under psychedelics.

Examining the correlation between the reported potency of the chosen psychedelic experience and how frequently different types of synthetic VH were selected, we found a significant positive correlation between potency and the iteration level preference (the slope value of the regression line of image selection frequency against iteration level), participants with highly potent psychedelic experiences were more likely to select psychedelic complex synthetic VHs with higher iteration levels (Pearson’s r = 0.33, p = 0.003, BF10 = 10.34). We found the same pattern of results for simple synthetic VHs: a significant positive correlation between potency and the iteration level preference for simple synthetic psychedelic VHs (Pearson’s r = 0.31, p = 0.007, BF10 = 5.344). These results suggest that manipulating the number of iterations used to produce synthetic VHs captures the visual characteristics of differing subjective intensities of psychedelic experience, with higher potency psychedelic experiences being better characterised by synthetic VHs produced using greater numbers of iterations.

Next, we assessed the veridicality of psychedelic complex VHs, by asking participants to rate how similar their VHs were to their normal visual experiences on a scale of 1 (identical, high veridicality) to 10 (completely different, low veridicality). Half the participants (38/77) reported that their psychedelic experience had included complex VHs and provided an average veridicality rating of 7.8 (SD=2.1) (Figure 13). In line with previous findings (Carhart-Harris et al., 2016; Preller & Vollenweider, 2018; Sanz et al., 2018; Studerus et al., 2011) these results suggest that for these participants their complex psychedelic VHs displayed relatively low veridicality.

**Figure 13.**
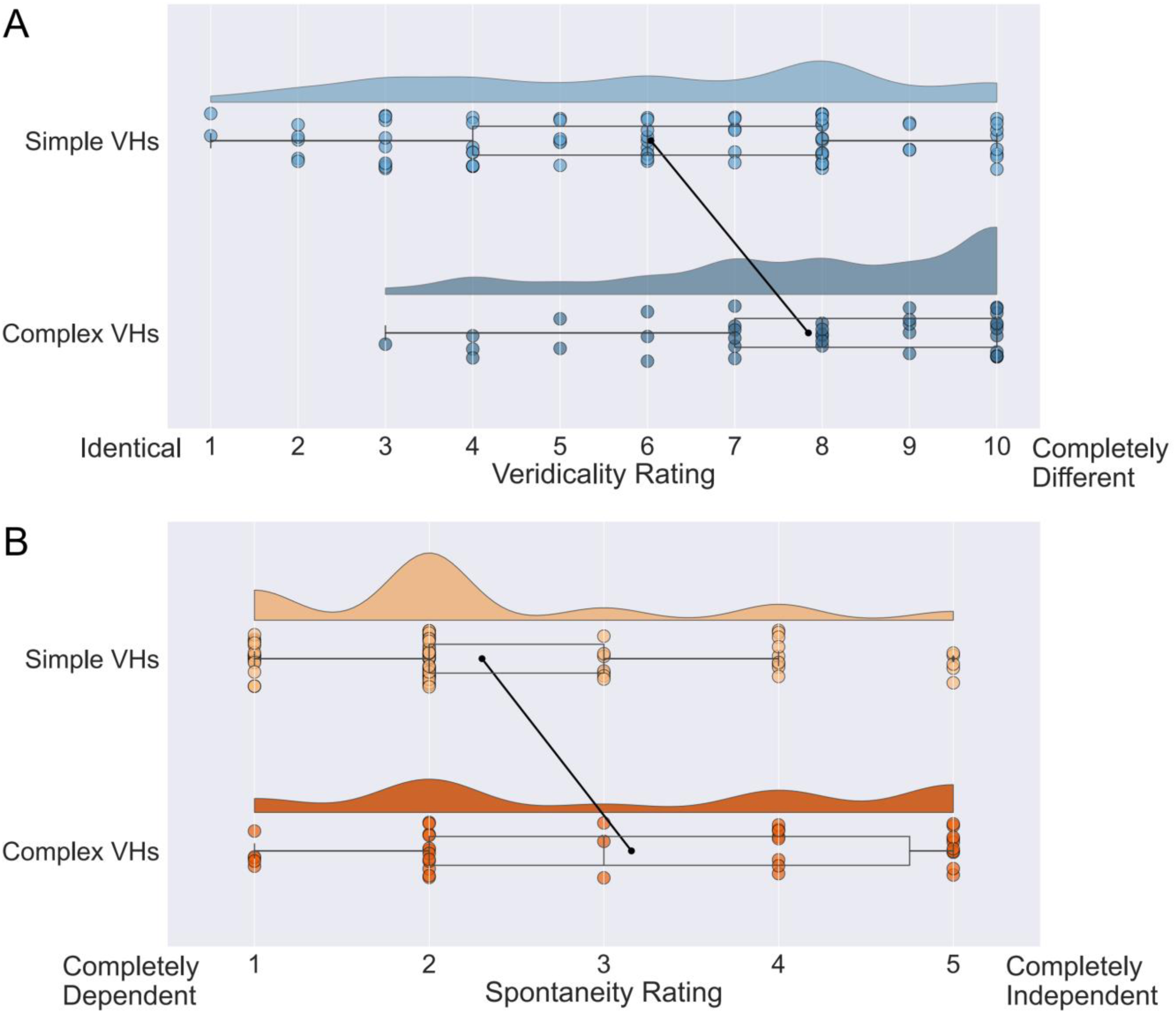
Raincloud plots displaying veridicality and spontaneity ratings for all participants in the Psychedelic Survey for both simple and complex psychedelic VHs. A. The veridicality rating displays the similarity of a participant’s chosen psychedelic experience (simple and complex) to their normal visual experience. B. Displays spontaneity ratings for all participants, reflecting how dependent their simple or complex VHs were on the existing content of a visual scene. Each circle indicates the data point for a single participant. Black lines between simple and complex plots indicate the mean values for each category.

We also asked the same participants to report the spontaneity of their complex VHs using a scale of 1 (completely dependent) to 5 (completely independent). We found an average spontaneity rating of 3.16 (SD=1.50) (Figure 13), suggesting that psychedelic complex VHs can be both dependent on, and independent of, sensory input, with a slight tendency to be rated more spontaneous in nature. We found no significant correlations between the potency of the chosen psychedelic experience and the reported veridicality (Pearson’s r = 0.27, p = 0.10, BF10 = 0.76) or spontaneity ratings (Pearson’s r = 0.19, p = 0.27, BF10 = 0.37).

Next, we assessed the veridicality and spontaneity of the reported simple VHs using the same method (Figure 13). All participants reported that they experienced simple VHs (one participant’s data was missing for these questions). We found an average rating for veridicality of 6.0 (SD=2.6), suggesting that the majority of psychedelic simple VHs were distinct from normal visual experiences of simple patterning (Figure 13). The average spontaneity rating was 2.3 (SD=1.1), suggesting that psychedelic simple VHs were more likely to be rated as transformations of existing features within the visual scene, than as spontaneous in nature. As with complex VHs, we found no significant correlations between the potency of the chosen psychedelic experience and ratings of veridicality (Pearson’s r = 0.22, p = 0.06, BF10 = 0.80) or spontaneity (Pearson’s r = 0.17, p = 0.15, BF10 = 0.40).

### 3.7 Comparing the veridicality of complex VHs between clinical and psychedelic groups

To explore the veridicality of complex VHs between psychedelic (n=38) and neurological/CBS (n=22) participants, we performed an exploratory analysis of the mean veridicality ratings between these groups. An independent sample t-test revealed that neurological/CBS participants rated the veridicality of their complex VHs significantly higher (M=8.3, SD=1.8) compared to the psychedelic participants (M=3.2, SD=2.1) (t(58)=9.505, p <0.001, Cohens’d=2.55, BF10=2.43e+10). Note that the veridicality rating scale for the psychedelic group was inverted to match the scale used in the clinical interview.

## 4 Discussion

Visual hallucinations offer fascinating insights into the mechanisms underlying perceptual experience, yet relatively little work has focused on understanding the differences in the phenomenology of VHs associated with different aetiologies. Using a computational neurophenomenological approach we first identified three dimensions of hallucinatory phenomenology which broadly characterise variations in VHs arising from neurological, CBS and psychedelic origins: their veridicality, spontaneity, and complexity. Using a coupled DCNN-DGN neural network architecture, we generated synthetic VHs that captured the differences in these three phenomenological dimensions between neurological, CBS, and psychedelic VHs. Each class of synthetic VH was generated by tuning the parameters of the visualisation algorithm to match the phenomenological dimensions represented in each group’s VHs. The parameters we manipulated were the inclusion or omission of the natural image prior of the DGN (corresponding to veridicality), altering the error function used to select the target neuron(s) within the DCNN (corresponding to spontaneity), and restricting the level within the DCNN that AM terminates (corresponding to complexity).

We verified the validity of this approach experimentally in two separate studies that investigated variations in hallucinatory experience in neurological-CBS patients and people with recent psychedelic experience. Both studies first verified that the three phenomenological dimensions usefully distinguished the different kinds of hallucination, and then asked whether the appropriate synthetic VHs were able to capture specific aspects of hallucinatory phenomenology for each aetiology. In both cases, we found that the relevant synthetic VHs were rated as being most representative of each group’s hallucinatory experience, compared to other synthetic VHs produced by the model. Together, our findings highlight how deep neural network architectures can be used to shed light on the computational mechanisms underpinning atypical perceptual phenomenology.

### 4.1 Simulating neurological and CBS VHs

Typically, neurological and CBS complex VHs are reported as being similar in visual quality to normal perception, that is, they display high veridicality (Fénelon et al., 2000; Frucht & Bernsohn, 2002; Mosimann et al., 2006; Papapetropoulos et al., 2008; Teunisse et al., 1996). The results of our phenomenological interview confirmed this aspect of hallucinatory phenomenology, with 92% of neurological and 90% of CBS participants rating their VHs as being ‘as real’ or ‘similar to’ their normal visual experiences (Table 3). We used DGN-AM to simulate this specific aspect of hallucinatory experience, finding that the learned natural image prior of the DGN resulted in synthetic VHs that displayed high veridicality (Figures 6 and 7), which was reflected by a comparable Inception Score to the benchmark simulation of non-hallucinatory perceptual phenomenology (Figure 5).

Complex neurological and CBS VHs are also reported to occur spontaneously – i.e., they are not transformations of content within a perceived scene (Frucht & Bernsohn, 2002; Mosimann et al., 2006; Papapetropoulos et al., 2008; Schultz & Melzack, 1991; Teunisse et al., 1996). Indeed, we found that 83% of neurological and 100% of CBS participants reported that their hallucinatory experiences occurred spontaneously (Table 3). We attempted to capture this phenomenological characteristic by using an error function for synthetic image generation that selects a target neuron independently of the input image (Fixed), entailing that the resulting synthetic VH was based on the categorical information represented by the target neuron and not the visual features of the input image.

In terms of the complexity of the hallucinatory experience, neurological VHs are most commonly reported as being complex in nature (Frucht & Bernsohn, 2002; Mosimann et al., 2006; Papapetropoulos et al., 2008). Our results again confirmed this aspect of hallucinatory phenomenology, finding that 83% of neurological participants reported that their VHs consisted of only complex VHs (see Table 2). We simulated this aspect of hallucinatory phenomenology by restricting the hierarchical level used to create the synthetic image; specifically, by ensuring that AM terminates to the highest categorical layer in the DCNN (Figure 2). In contrast, the most commonly reported class of VHs reported in CBS are simple in nature (Abbott et al., 2007; ffytche & Howard, 1999; Santhouse et al., 2000; Schultz & Melzack, 1991; Teunisse et al., 1996). We found that 90% of CBS participants reported not only simple but also complex VHs. Restricting the AM termination level within the DCNN to a lower layer (conv4) resulted in synthetic VHs that displayed many indicative features of simple CBS VHs (Figure 7). This was due to the overemphasis of the visual features learnt by neurons in this layer during training, together with the influence of the learned natural image prior from the DGN, i.e., low-level colours and textures associated with natural real-world objects.

Verifying the validity of our synthetic VHs, we found that both neurological and CBS participants reported that they provided a close approximation of both their simple and complex hallucinatory experiences (Figures 6 and 7). In an exploratory analysis, we examined which iteration value used to generate the synthetic VHs participants selected as being most representative of their complex VHs, where iteration value roughly translates to the degree to which the input image is altered. Here, we found that, on average, neurological participants chose synthetic VHs with a low number of iterations (average iteration value of 3; see Figure 3). In contrast, CBS participants, on average, selected a higher iteration value (average iteration value of 18.6) as being the most representative of their complex VHs. Together these findings fit the intuition that for these groups, neurological participants experience hallucinatory phenomenology that is close to, but not identical to, their normal perceptual experience, while CBS participants have complex hallucinatory experiences with less veridicality.

Summarising the performance of our model in simulating neurological and CBS VHs, we found that the addition of the natural image prior of the DGN, combined with the Fixed error function, and the hierarchical depth at which AM terminates, allowed us to simulate three specific aspects of neurological and CBS hallucinatory phenomenology: their veridicality, spontaneity and complexity. The results of the semi-structured interviews with these patients verified that these properties were distinct aspects of the perceptual phenomenology associated with neurological and CBS VHs and that our models’ synthetic VHs provided representative examples of both simple and complex hallucinatory experience in these groups.

### 4.2 Simulating psychedelic VHs

In contrast to complex neurological or CBS VHs, psychedelic VHs are typically reported as having reduced veridicality, due to their dream-like qualities and the inclusion of visual distortion and unrealistic colours and patterning (Carhart-Harris et al., 2016; Kometer et al., 2013; Kometer & Vollenweider, 2018; Preller & Vollenweider, 2018; Sanz et al., 2018; Studerus et al., 2011; Timmermann et al., 2018).

To simulate the diminished veridicality associated with psychedelic VHs, we suppressed the influence of the learned natural image prior in the generation of synthetic VHs by removing the contribution of the DGN from the model (Figure 8). This resulted in the use of a visualisation method (DCNN-AM) that was identical to our previous simulations of psychedelic hallucinatory phenomenology (Suzuki et al., 2017). The resulting synthetic VHs displayed striking ‘hallucinatory’ qualities, bearing intuitive similarities to a wide range of psychedelic VHs reported in the literature (Carhart-Harris et al., 2016; Kometer et al., 2013; Kometer & Vollenweider, 2018; Preller & Vollenweider, 2018; Studerus et al., 2011; Timmermann et al., 2018).

The results of the online psychedelic survey confirmed the diminished veridicality associated with psychedelic VHs, with participants rating their complex VHs as displaying low veridicality (Figure 13). This finding was further supported by an exploratory analysis comparing the veridicality of complex VHs between psychedelic and neurological/CBS groups, which found that complex psychedelic VHs were rated as displaying significantly lower veridicality than both neurological and CBS VHs (see section 3.7). The reduction in veridicality of this class of synthetic VH (Figure 8) was also reflected by a substantially reduced Inception Score compared to both benchmark and simulations of neurological and CBS complex VHs (Table 2).

Complex psychedelic VHs are also commonly reported as being less spontaneous than neurological/CBS VHs, often appearing to be transformations of existing information within a visual scene (Kometer & Vollenweider, 2018; Preller & Vollenweider, 2018; Swanson, 2018). We simulated this aspect of hallucinatory phenomenology by using an error function that selected target neurons within the DCNN based on the visual features of the input image (Deep-Dream). However, the results of the online psychedelic survey revealed considerable heterogeneity in this aspect of hallucinatory phenomenology (Figure 13). Psychedelic complex VHs were rated as being both dependent or independent of sensory input, with a slight tendency to be rated more frequently as spontaneous in nature. In contrast, simple VHs were more likely to be rated as not being spontaneous, i.e., as being transformations of existing visual input. One possible explanation for this variability is that there may be a confound of dose-dependency, with both complexity and spontaneity of VHs increasing with dose. This would be in line with findings of the prevalence of complex hallucinations increasing with drug dose (Shulgin, 1997; Strassman et al., 1994; Shanon, 2002; Strassmann, 2001; Studerus et al., 2011; Leptourgos et al., 2020). Examining the number of iterations used to produce synthetic psychedelic VHs, we found a significant positive correlation between iteration number (of selected images) and the potency of psychedelic taken, suggesting that, not only did our synthetic VHs closely resemble psychedelic experience, but synthetic VHs produced using differing numbers of iterations was able to capture variations in the intensity of the visual phenomenology of psychedelic experience.

Psychedelic VHs occur in both complex and simple forms, as with simple CBS VHs. We simulated simple psychedelic VHs by restricting the hierarchical level within the DCNN to which information propagates (i.e., the level at which AM terminates) to a lower layer (conv4) (Figure 9). Compared to simulations of simple CBS VHs (Figure 7), synthetic psychedelic VHs included a high prevalence of geometric shapes, colours and patterning, resembling typical reports of psychedelic simple VHs (Bressloff et al., 2001; Cowan, 2015; Díaz, 2010; Nichols, 2016; Tyler, 1978).

Verifying the validity of psychedelic synthetic VHs, using an online forced choice image selection task, we found that participants who reported complex VHs chose psychedelic synthetic VHs significantly more frequently than synthetic neurological VHs as being representative of their experience. Similarly, participants chose simple psychedelic synthetic VHs significantly more frequently than synthetic CBS VHs (Figure 11) as most closely resembling their psychedelic experience. Out of participants who indicated they had experienced simple VHs (n=81), we found that on average they were most likely to choose synthetic VHs generated using a low number of iterations (10) (see Figure 12), suggesting that for this sample their simple hallucinatory phenomenology was relatively subtle in nature, consisting of low-level visual distortion, rather than the colourful kaleidoscopic patterning usually associated with classical hallucinogens.

Summarising the performance of our model in simulating simple and complex psychedelic VHs, we found that suppressing the natural image prior (DGN), combined with the Deep-Dream error function, and constraining the hierarchical depth at which AM terminates, produced synthetic VHs that resemble many aspects of psychedelic hallucinatory phenomenology, including their reduced veridicality, spontaneity, and differing levels of complexity. The results of the image selection task verified that these synthetic VHs provided a representative depiction of both simple and complex psychedelic hallucinatory experience. They also support the conclusion that psychedelic complex VHs display significantly reduced veridicality compared to neurological or CBS VHs, and revealed that, unlike neurological or CBS VHs, complex psychedelic VHs could be experienced as being both spontaneous or dependent on sensory input.

### 4.3 Predictive processing accounts of hallucinatory experience

By decoupling perceptual content from normal sources of sensory input, VHs exemplify the constructive nature of perceptual experience. They demonstrate that the brain is capable of generating rich, detailed and in some cases life-like perceptions irrespective of (or only partly constrained by) actual sensory input (dreams, of course, show this too). It is therefore natural to interpret VHs within ‘predictive processing’ (PP) accounts of perception and brain function (Rao & Ballard, 1999; Friston 2005; Clark, 2013; Seth, 2014).

Indeed, considerable work has focused on understanding the computational origins of VHs within a PP framework. Such studies generally associate VHs – neurological and clinical VHs in particular – with aberrant perceptual inference, usually appealing to overly strong perceptual priors (Corlett et al., 2019; Friston, 2005; O’Callaghan et al., 2017; Powers et al., 2017). For example, O’Callaghan et al., (2017) found that in PD, patients who experienced VHs displayed slow and inefficient sensory processing in a perceptual decision-making task. They interpreted this result within a Bayesian framework, suggesting that if visual information is accumulated slowly and inefficiently it will be deemed less informative, and may be down-weighted leading to an over-reliance on top-down expectations, resulting in the experience of VHs.

While we acknowledge that the model used in this paper is not an explicit PP model, the architecture and visualisation methods used here display many similarities to PP models in terms of hierarchical information processing. For example, the DCNN used here is a purely feedforward network, which we employed to generate synthetic VHs using both DCNN-AM and DGN-AM. Both instances of AM optimise the input image via backpropagation, which at least informally, mirrors certain aspects of the top-down signalling that is central to PP accounts of perception [see for example (Millidge et al., 2020), who show that predictive coding networks can implement an approximation to backpropagation].

Specific parallels to PP can also be identified in the parameters used to generate each type of synthetic VH. We simulated neurological/CBS VHs using the Fixed error function and DGN-AM. The Fixed error function resulted in the input image being overwritten by synthetic hallucinatory content, based on the features represented by a single target neuron. From a PP perspective, this process bears intuitive similarities to theories that posit that this type of VHs occurs due to overly strong perceptual priors overwhelming sensory prediction error signals (O’Callaghan et al., 2017; Powers et al., 2016).

Interpreting the contribution of the DGN within a PP framework, we emphasise that the natural image prior of the DGN should not be confused with perceptual priors. Instead, it can be thought of as a variety of hyperprior, which results in each layer and neuron of the network having a tendency to generate realistic interpretable images. Other examples of hyperpriors within a PP system include the physical constraints imposed by space and time e.g., “that there is only one object (one cause of sensory input) in one place, at a given scale, at a given moment,” (Clark, 2013, p. 196). Such spatial and temporal hyperpriors narrow and restrict the range of possible hypotheses about the external objects causing incoming sensory input (Clark, 2013), increasing the tractability of (approximate) inference (Swanson, 2016; Tenenbaum et al., 2011; Blokpoel et al., 2012; Clark, 2013; Kwisthout, 2014; Baltieri, 2020). Similarly, the hyperprior of the DGN constrains the vast set of possible uninterpretable images that may excite a target neuron, forcing the network towards producing synthetic outputs that are both interpretable and that closely resemble natural images.

In contrast to clinical VHs, convergent evidence has suggested that psychedelic VHs occur due to a reduction in top-down control and an increase in bottom-up information transfer (Alonso et al., 2015; Carhart-Harris & Friston, 2019; Muthukumaraswamy et al., 2013; Schartner & Timmermann, 2020; Timmermann et al., 2018). Such relaxation of high-level priors has been proposed to allow ascending prediction errors from lower levels of the visual system to reach higher-than-normal levels of the visual hierarchy and be integrated into conscious experience (Carhart-Harris & Friston, 2019). Our use of the Deep-Dream error function and DCNN-AM to generate synthetic psychedelic VHs intuitively approximates this disruption in the balance between hierarchical information processing. From a PP perspective, compared to the Fixed or WTA, the Deep-Dream error function has the effect of relaxing the perceptual priors imposed on incoming sensory data. The selection of multiple target neurons based on the visual characteristics of the input image by this error function lowers the precision of the perceptual prior, effectively increasing bottom-up information transfer.

### 4.4 Alternative machine learning models for computational neurophenomenology

The coupled DGN-DCNN network we used served to capture some of the core features of a PP architecture in a way well suited to synthetic image generation. While PP architectures such as predictive coding (PC) networks offer a model of cortical function that exhibits many similarities to our present understanding of human visual perception (Rao & Ballard, 1999; Friston 2005), current limitations of this class of model, such as the scalability of local learning rules (Bartunov et al., 2018), their restriction to low-dimensional images and narrowly focused datasets, preclude their use in simulating visual phenomenology. In contrast, machine learning models such as DCNNs are highly scalable and, when coupled with DGNs, have the potential to shed new light on the neural mechanisms underlying visual phenomenology in a way that reflects key principles of theories of predictive perception.

DCNNs have long been utilised as models of the human visual system. Their utility is exemplified by close correspondences between activity within layers of DCNNs and activity in regions along the ventral visual stream. For example, a comparison of internal representational structures of trained DCNNs and the human visual system performing similar object recognition tasks has revealed similarities between the representational spaces in these two distinct systems (Cadieu et al., 2014; Grossman et al., 2019; Khaligh-Razavi & Kriegeskorte, 2014). However, others have argued that the correspondences between brain and model activity involve shared, not task-relevant, variance and more rigorous tests are required to provide a stricter measure of the correspondence between these two systems (Sexton et al., 2022).

More fundamentally, the prevalence of top-down connectivity in the human visual system points to a fundamental difference with DCNNs, which implement top-down influence purely through the learning algorithm (typically backpropagation). A productive direction for future research would therefore be to address the current limitations in using PP-like architectures – such as predictive coding (PC) – for generating simulated phenomenology.

One promising possibility is raised by work demonstrating certain functional equivalences between backpropagation and predictive processing (Whittington & Bogacz, 2017; Millidge, Tschantz, & Buckley, 2020). Here, the bottom-up classification process employed by DCNNs are viewed as predictions from data to labels, and the error backpropagation that refines the network weights are viewed as ‘prediction errors’. Based on this work, a recent ‘hybrid predictive coding’ network architecture has been proposed (Tschantz, et al., 2022) in which predictions and prediction errors flow bidirectionally in hierarchical networks, to adaptively balance ‘fast’ and ‘slow’ inference. Hybrid predictive coding networks combine ‘amortised inference’ – which performs ‘fast and cheap’ approximate perceptual inference for familiar data – with the iterative inference implemented in standard predictive coding. By combining fast and slow inference, hybrid predictive coding networks are well suited for shedding light on the biological relevance of the feedforward sweeps and iterative recurrent activity observed in the visual cortex during perceptual tasks. Future work could also examine how adaptations of hybrid predictive coding could be used to provide insights into aspects of hallucinatory phenomenology.

### 4.5 Limitations

There are several limitations worth highlighting in the current investigation. First, the benchmark performance of the model did not succeed in producing output images that were identical to the input images. This finding may have been due, in part, to the synthetic images generated by the DGN being based on the compressed latent vectors within higher layers of the DGN and not the pixel values of the input image. The compression of the features of the input image has the effect of diminishing the realism of the output image. Another contributing factor is likely to be the number of separate categories the DCNN has been trained to classify, which while relatively large (1,000 classes), is trivial compared to the human brain. This constraint means that when presented with an arbitrary (untrained) input image the model cannot generate an output that displays a fine-grained visual similarity to the input image. Using larger training-set data would ameliorate this problem.

Another important limitation that should be considered when interpreting these results is the relatively small sample sizes in both studies investigating the phenomenology of different hallucinatory experiences. We acknowledge the conclusions drawn here are limited to this specific groups of participants. Further work is required to establish if the phenomenological differences in VHs found here, both across and within different aetiological categories, apply to wider populations.

In an attempt to assess how representative our synthetic VHs were of aetiologically distinct hallucinatory experiences we employed different methodological approaches, we enquired about the general phenomenology of a person’s VHs, including specific questions about the complexity, veridicality, and spontaneity of their VHs. Our model also allowed us to ask participants to directly rate the visual similarity between the appropriate synthetic VHs and their hallucinatory experiences. The flexibility of our model allowed us to create a wide range of images produced using differing number of iterations or constraining the hierarchical depth at which AM terminated. While we highlight the success of this approach, we also acknowledge the inherent challenges in assessing how closely an individual’s private subjective experience is captured by a synthetic VHs.

Finally, as discussed above, there is increasing evidence highlighting the lack of biological plausibility of the architecture and learning rules used by DCNNs. Future studies attempting to generate simulations of phenomenology should consider using PP-like architectures.

## 5 Conclusion

We have described a novel method for simulating altered visual phenomenology associated with neurological, CBS and psychedelic VHs. Using a coupled deep convolutional neural network (DCNN) and deep generator network (DGN) we leveraged recent advances in visualising the learned representations of these networks to simulate three dimensions relevant to distinguishing aetiologically distinct VHs: their realism (veridicality), their dependence on sensory input (spontaneity), and their complexity. By selectively manipulating the parameters of the algorithm used to generate the synthetic VHs, we were able to capture differences in these phenomenological characteristics across distinct VH populations. Two experimental studies confirmed the existence of variations in these phenomenological dimensions across groups and critically, that the appropriate synthetic VHs were representative of the hallucinatory phenomenology experienced by neurological- CBS and psychedelic groups.

Overall, our findings highlight the phenomenological diversity of VHs associated with distinct causal factors, and the need for a more fine-grained mapping between computational mechanisms and specific hallucinatory phenomenology. We have shown here how a coupled machine learning network and specific visualisation algorithms can be used as a step towards this goal, by demonstrating that such models can successfully capture the distinctive visual characteristics of hallucinatory experience. Future research could usefully apply this approach using more biologically plausible predictive processing architectures to test computational hypotheses about the origins of specific hallucinatory phenomenology and validate their results within appropriate hallucinatory populations. In this way, it would be possible to further close the loop between hallucinatory experience and their computational mechanisms.

## 6 Conflict of Interest

The authors declare that the research was conducted in the absence of any commercial or financial relationships that could be construed as a potential conflict of interest.

## 7 Author Contributions

K.S. conceived the design of the study. K.S. and D.J.S wrote the manuscript. All authors contributed to revising this manuscript at all stages.

## 8 Funding

All authors are grateful to the Dr Mortimer and Theresa Sackler Foundation, which supports the Sussex Centre for Consciousness Science. A.K.S. acknowledges additional support from the Canadian Institute for Advanced Research (CIFAR) via their Azrieli Programme on Brain, Mind, and Consciousness.

## Acknowledgments

We would like to thank Sarah Wolffe for assistance with data collection and Alexander Tschantz and Beren Millidge for helpful discussions. We would also like to thank Parkinson’s UK and Esme’s Umbrella for their invaluable assistance in helping recruit participants for this study.

## Glossary

Activation Maximisation (DCNN-AM): A method which maximises the activity of a particular neuron (group of neurons or an entire layer) within a (D)CNN by adjusting the network’s input (e.g., through backpropagation), while keeping the weights and outputs of the network fixed. AM is a widely used approach to visualise the representations learned by a specific neuron (or neurons) within a (D)CNN. Because the form of AM that we use is restricted to a DCNN we refer to it as DCNN-AM.
Backpropagation: Short for ℌbackward propagation of errors”; backpropagation (or backprop) is a widely used algorithm for the training of artificial neural networks using gradient descent. Given a network and an error function, it enables the calculation of the gradient of the error function with respect to the weights in the network. Each weight is then adjusted in proportion to how much it contributes to the overall error so that the network improves its performance over successive iterations.
Computational (neuro)phenomenology: We use computational phenomenology to describe the use of computational modelling to model phenomenological properties, rather than (or in addition to) function or behaviour. This approach becomes computational neurophenomenology when the computational models can be related to neural mechanisms, either by interpretation or through generating experimental predictions (Suzuki et al 2017, Ramstead et al 2022).
(Deep) Convolutional Neural Network ((D)CNN): A class of (deep) neural networks with an architecture specifically designed to process pixel-wise data of two-dimensional images. DCNNs are trained via backpropagation to classify the content of a set of training images into distinct categories, creating a feature map that summarises the presence of the detected features within an input image. DCNNs have been used primarily for applications such as object detection and image classification.
Deep Generator Network (DGN): A class of deep neural network that is used to generate synthetic data. In this paper, we use the term DGN to refer to a specific type of deep generator network that has been pre-trained using the generative adversarial network (GAN) framework.
Deep Generator Network Activation Maximisation (DGN-AM): A novel form of AM proposed by Nguyen et al. (2016). Instead of directly optimising the image to maximise the activity of a target neuron within a DCNN as in DCNN-AM, Nguyen et al. (2016) used the learned natural image prior of a pre-trained DGN in combination with DCNN-AM to synthesise a new image that maximised activity in a target neuron within a DCNN, resulting in realistic and interpretable output images.
Error Function: A method used in machine learning to minimise the error between a predicted value and an actual value, which is commonly used to evaluate the performance of the model. For example, in supervised learning for image recognition, an error function calculates the discrepancy between the actual label of an image, and the model’s estimation of this value. The network then attempts to minimise the error for each iteration using (for example) backpropagation.
Generative Adversarial Network (GAN): The GAN is a learning framework in which a ‘generator’ and ‘discriminator’ network are pitted against each other in an adversarial manner. A generative neural network is used to generate synthetic data (e.g., images) that resemble a training data set. The discriminator network (e.g., a DCNN) is then trained to distinguish images produced by the DGN from those in the actual training data. The two networks are trained together in an adversarial zero-sum game in which the DGN tries to ‘fool’ the discriminator, while the discriminator tries to keep from being fooled. GANs are widely used in image processing to generate realistic synthetic images.
Gradient descent: A popular class of optimization algorithms in which the local minimum of a function is found by iteratively moving in the direction of the steepest descent as defined by the negative of the gradient. In machine learning, gradient descent is often used to update the parameters (weights) of a neural network, as for example in backpropagation.
Inception Score (IS): A widely used objective approach to assess the realism of synthetic images produced by generative models, such as a GAN (Salimans et al., 2016).
Natural Image Prior: A common method used to constrain optimization of a (D)CNN towards generating synthetic images that resemble natural images, for example by minimising the colour difference between adjacent pixels (hand-designed natural image prior). Recently, Nguyen et al., (2016) used the learned natural image priors acquired by a DGN during GAN to constrain the output of the DCNN to generate synthetic images that closely resembled natural images.

## Notes

https://osf.io/nr4ke/files/osfstorage

### Competing Interest Statement

The authors have declared no competing interest.

