## Supplementary Material for "Modelling Phenomenological Differences in Aetiologically Distinct Visual Hallucinations Using Deep Neural Networks"

Modelling Phenomenological Differences in

Aetiologically Distinct Visual Hallucinations

Using Deep Neural Networks

**Keisuke Suzuki*, David J. Schwartzman & Anil K. Seth**

### 1 Supplemental Methods

#### 1.1 Model Architecture and Parameters

##### 1.1.1 Pre-training of DGN

The DGN used by Nguyen et al., (2016) was taken from Dosovitskiy & Brox, (2016a) and was pre-trained using the GAN framework, combined with the representation learning method (Figure 1). This DGN consists of nine up-convolutional (up-sampling and a subsequent convolutional) layers (u1-u9) and three fully connected layers (fc5, fc6, fc7) (Figure S1) which are designed to invert a convolutional network. The variant of GAN training here involved three networks: ***C***, a DCNN (CaffeNet), ***G*,** a DGN, and ***D***, a discriminator network. ***G*** learns to synthesise images that are similar to the training data set, while ***D*** learns to distinguish the images produced by ***G*** from the actual training data set. Formally, the GAN framework minimises the following loss functions (*L_discr_* and *L_adv_*) for ***D*** and ***G***, respectively.

$L_{discr}= -\sum_{i} \log\left( D\left( x_{i} \right) \right) +\log\left( 1-D\left( G\left( y_{i} \right) \right) \right)$

$L_{adv}= -\sum_{i} log(D(G(y_{i})))$

where ***x*** denotes the training image and ***y*** denotes the latent value of ***G***.

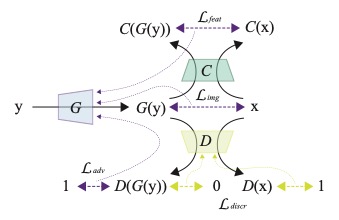

**Figure S1.** The GAN framework used in Dosovitskiy & Brox, (2016). The model uses three different loss functions: feature loss (L_feat_), image loss (L_img_), and adversarial loss (L_adv_). Black solid lines denote the forward pass. Dashed, purple and yellow lines with arrows on both ends denote the losses. Thin dashed lines denote the flow of gradients. The image is adapted with permission from Dosovitskiy & Brox, (2016).

In the model from Dosovitskiy & Brox (2016a), in addition to minimising the adversarial losses, ***G*** is also trained to minimise two additional losses. The first loss function (*L_img_*) attempts to minimise the difference in the images generated by ***G*** to the training images at the pixel level. This is formally described as minimising the image loss (*L_img_*):

$L_{img}= \sum_{i} \left. \left\| G\left( y_{i} \right)- x_{i} \right\| \right._{2}^{2}$

where ||...||^2^_2_ indicates the L2 normalisation. The second loss function (*L_feat_*) attempts to minimise the mismatch between the categorical representation of ***C*** based on the images generated by ***G*** (i.e., the feature vector at the top layer of ***C***) and the categorical representation of ***C*** based on the training image. This is formally described as minimising the feature loss (*L_feat_*) as follows:

$L_{feat}= \sum_{i} \left. \left\| \left. C\left( G\left( y_{i} \right) \right)- C\left( x_{i} \right) \right. \right\| \right._{2}^{2}$

After training minimises all the above losses ***G*** is capable of generating ‘realistic’ images based on a random latent value from one of the fully connected layers (fc6 in Figure 1) as an inversion of the DCNN. We used ***C*** and ***G*** in this study, ***D*** was only used in the training of these networks by Dosovitskiy & Brox (2016a) and was not used in this study.

##### 1.1.2 Synthesising preferred images using AM

**DCNN-AM**

This optimisation process performed by DCNN-AM which tries to find **x** to maximise the target neuron ***h*** in the DCNN ***C*** can formally be described as:

$$\hat{x}=\underset{x}{\mathrm{argmax}} \left( C_{h}\left( x \right)-\lambda\left| \left| x \right| \right| \right)$$

where λ ||x|| is a regularisation term. $\lambda$ = 0.005 is taken from the original study (Nguyen et al., 2016).

**DGN-AM**

Formally, DGN-AM searches the latent space of ***G*** to find a value ***y*** such that ***G(y)*** is an image that maximises activation of the target neuron ***h*** in the DCNN ***C***. This process can be formalised as finding $\hat{y}$ in the following equation:

$$\hat{y}=\underset{y}{\mathrm{argmax}} \left( C_{h}\left( G\left( y \right) \right)-\lambda\left| \left| y \right| \right| \right)$$

with a regularisation term ($\lambda$ = 0.005).

##### 1.1.3 Pseudo code used to generate synthetic VHs

```
if ( withDGN ) // DGN-AM
x = get_initial_image_through_DGN(x_input) // (1)
else
x = x_input
end

for ( the_number_of_iterations )
v = DCNN_forward_path(x) // (2)terminates at fc8 or conv3/conv4
	v_AM = act_max(v) // (3) fixed or winner-take-all
	x_update = DCNN_backward_path(v_AM) // (4)

if ( withDGN ) // DGN-AM
x_diff = x - x_update // (5)
z_diff = DGN_backward_path(x_diff)
	z = z + z_diff // update the latent vector // (6)
x = DGN_forward_path(z) // (7)
else // DCNN-AM
x = x_update
end
end
x_output = x
```

**Table S1.** Pseudo code used to generate images. A variable, x, denotes the image data, v denotes the feature vector in the DCNN, z denotes the latent vector in the DGN. For DGN-AM, the process starts with an initialisation stage, where an image is obtained from the feature representation of the input image through the DGN (‘get_initial_image_through_DGN()’). The forward_path() and backward_path() are the functions used to process information through the networks in the opposite directions. DGN-AM (used in Nguyen et al., (2016)) is executed when ‘withDGN’ variable is true, whereas DCNN-AM is executed when ‘withDGN’ is false. For the DCNN-AM, the forward pass terminates at a pre-specified target layer (fc8, conv3, or conv4). The function ‘act_max()’ operates by maximising a target neuron depending on which type of AM was used.

##### 1.1.4 Additional Simulation Parameters

For all simulations, we used the same five arbitrary input images that were chosen as being representative of 5 categories (among 1,000) used in the training of the DCNN (bird, mushroom, lamp, volcano, and flower). The category of input images was randomly selected from the 1,000 unique categories of CaffeNet. We then performed an internet search to find a single exemplar of each category, using the specific CaffeNet label of each category (junco, bolete, lampshade, volcano, cardoon) and converted them to the same image size as CaffeNet images (227 x 227 pixels). This meant that all input images used in our simulations were unlikely to be part of the original CaffeNet training data set, making the results of our simulations more generalizable.

For all simulations using DCNN-AM we applied several minor optimisations. To reduce known artefacts in image generation when using AM (Olah et al., 2017), for every iteration we added a spatial jitter (-4 to +5 pixels for x and y respectively) and rotated the image (randomly chosen between -5 to +5 degrees). In addition, due to the absence of the DGN, the initialisation stage was omitted from the model: instead, an input image was passed directly to the DCNN.

For all simulations we ran the model for a total of 1,000 iterations. This value was derived from experimentation with the optimal number of iterations to allow DGN-AM to converge on a stable output. After extensive piloting, we found that 1,000 iterations were sufficient to ensure the output image was not altered any further.

##### 1.1.5 Images used to calculate Inception Scores

We used the Inception Score (IS)(Salimans et al., 2016) to provide an objective method of measuring the realism of our simulations of hallucinatory phenomenology. To calculate the IS an image is entered into an “inception network”, a class of DCNN (e.g., GoogleNet), trained for image classification (Szegedy et al., 2014). To calculate the IS we used the same 5 images as in all simulations, in addition, we selected 27 random image categories from CaffeNet and used the same approach outlined in section 2.1.5 to select 27 random images (see Figure S1 for the images used). An image produces the highest IS when it activates only a single neuron in the categorical layer of the DCNN, meaning that the features of the image converge into a single categorical label. We compared the IS between input images, benchmark, and all simulations of synthetic VHs.

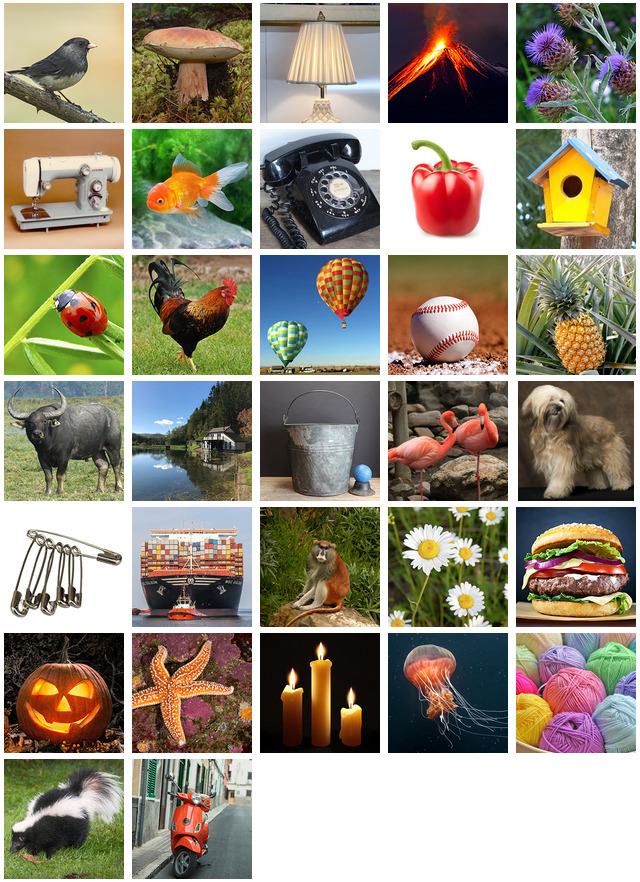

**Figure S1.** Unaltered arbitrary input image set used to calculate Inception Score, including the 5 images used in all simulations. Note that while all the images are representative of a specific trained category within the DCNN, they were not used in the training of the DGN or DCNN. This image set was used as the input images for all other simulations where Inception Scores are calculated.

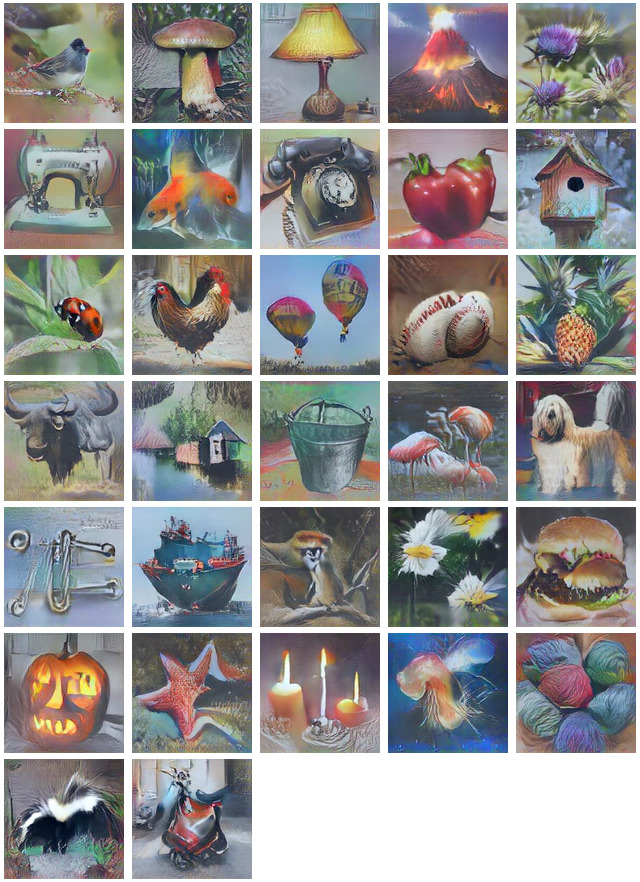

**Figure S2.** Outputs of DGN-AM following 1000 iterations, using winner-take-all error function simulating non-hallucinatory perceptual phenomenology. The resulting Inception Score was used as the benchmark of the model's performance in producing realistic synthetic images, which all other simulations were compared to.

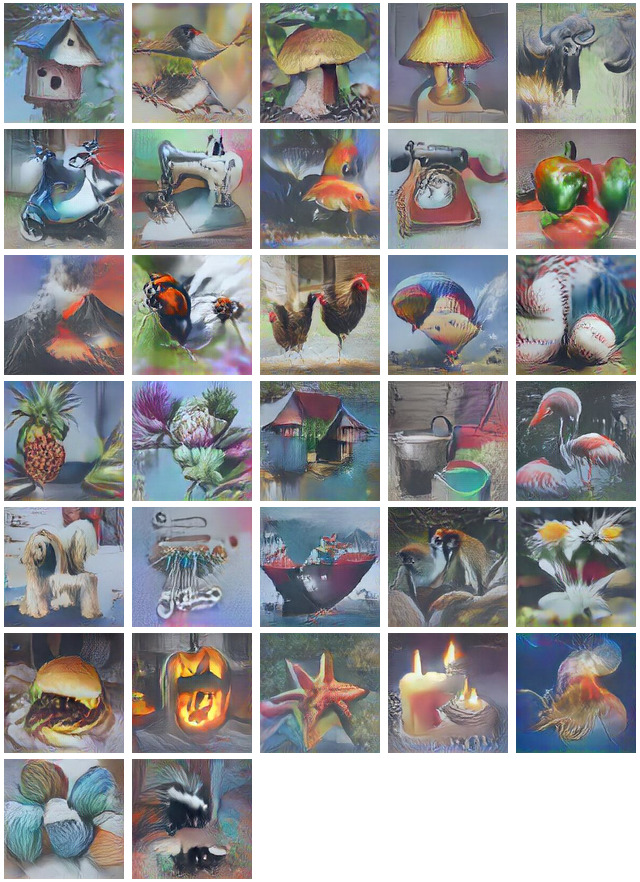

**Figure S3.** Outputs of DGN-AM following 1000 iterations, using the Fixed error function, simulating neurological complex VHs used to calculate Inception Scores. The predetermined categorical target neurons were from left to right: birdhouse, junco, bolete, lightshade, water buffalo, motor scooter, sewing machine, goldfish dial phone, bell pepper, volcano, ladybug, cock, balloons, baseball, pineapple, cardoon, boat house, buckets, flamingo, Tibetan terrier, safety pin, containership, patas monkey, daisy, hamburger, jack-o-lantern, starfish, candle, jellyfish, wool, skunk.

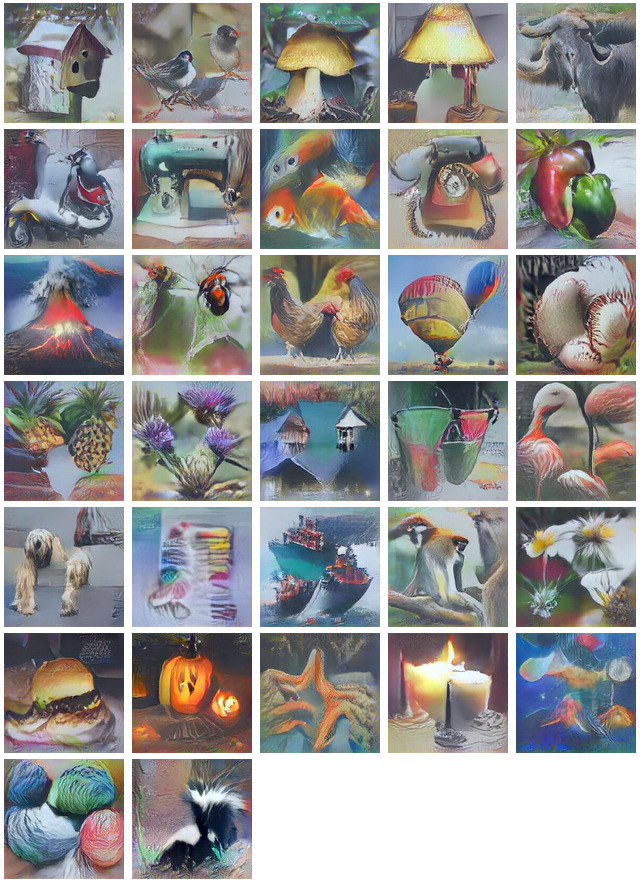

**Figure S4.** Outputs of DGN-AM using Fixed error function, simulating CBS Complex VHs used to calculate Inception Scores. The predetermined categorical target neurons were from left to right: birdhouse, junco, bolete, lightshade, water buffalo, motor scooter, sewing machine, goldfish dial phone, bell pepper, volcano, ladybug, cock, balloons, baseball, pineapple, cardoon, boat house, buckets, flamingo, Tibetan terrier, safety pin, containership, patas monkey, daisy, hamburger, jack-o-lantern, starfish, candle, jellyfish, wool, skunk.

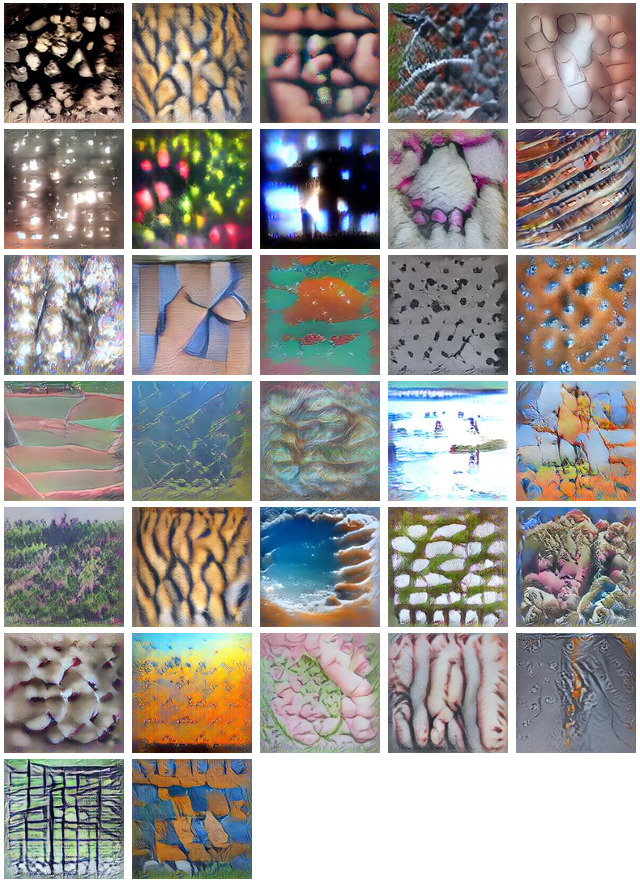

**Figure** **S5.** Outputs of DGN-AM following 1000 iterations, using the Fixed error function, simulating CBS simple VHs used to calculate Inception Scores. Activity was restricted to a lower layer of the DCNN (Conv4). The predetermined categorical target neuron for each simulation were randomly selected from the Conv4 layer.

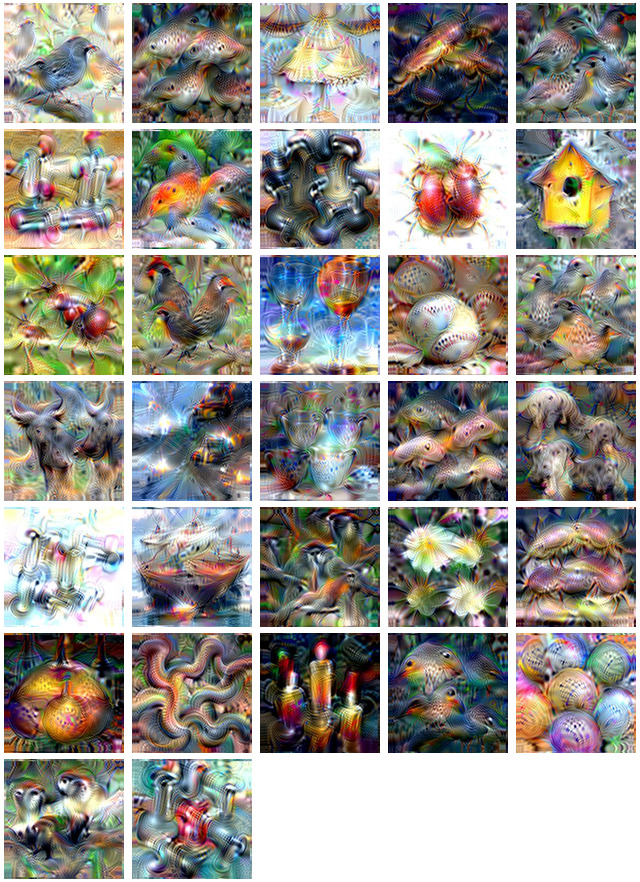

**Figure S6.** Outputs of DCCN-AM following 1000 iterations, using Deep-Dream error function, simulating psychedelic complex VHs used to calculate Inception Scores. Using Deep-Dream Error Function the target neurons within the highest layer of the DCNN that responded maximally to the input image were selected.

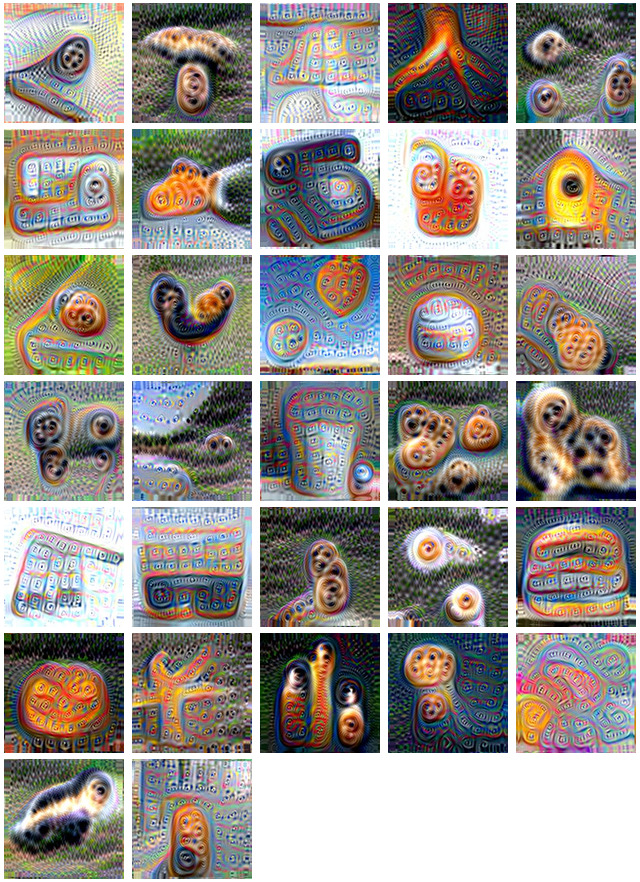

**Figure S7.** Outputs of DCCN-AM following 1000 iterations, using Deep-Dream error function, simulating psychedelic simple VHs used to calculate Inception Scores. Activity was restricted to a lower layer of the DCNN (Conv4). Using the Deep-Dream error function the target neurons within the Conv4 layer of the DCNN that responded maximally to the input image were selected.

#### 1.2 Psychedelic Survey Supplemental Methods

**Participants**

Participants were recruited through the Brighton Psychedelic Society, the Psychedelic Research Society of the University of Sussex, the UK Psychedelic Society and via social media advertising. Participants were asked to confirm that they were proficient in English language, had normal to corrected vision, had no previous traumatic psychedelic experiences, had no neurological or psychiatric conditions within the past year, and that they were not currently under the influence of drugs or alcohol. No identifying information was collected. Participants who completed the study were entered into a prize draw to win a £50 Amazon voucher.

**Materials**

The image set used in the survey was created by selecting 32 random image categories from CaffeNet and using the same approach outlined in section 2.1.5 to select random image categories and then find, via an internet search, exemplar images of each category (see <https://osf.io/nr4ke/files/osfstorage> for actual images used).

For the survey, we generated examples of synthetic VHs for each image representing four main categories: 1. Simple neurological, 2. Simple psychedelic, 3. Complex neurological, 4. Complex psychedelic. This process resulted in 32 synthetic VHs for each category. Each category of synthetic VH was generated using the model parameters described in section 2.1.4.

To investigate if the subjective potency of the psychedelic experience (see Procedure, below) correlated with the number of iterations an input image was processed by, we also investigated the effects of iteration level on image selection, by producing synthetic VHs for all four categories following 10, 100, and 1000 iterations.

For complex neurological and psychedelic categories, this led to a total image set of 384 images (input images x32, AM type x 2 (DGN-AM or DCNN-AM), error function x2 (WTA or Fixed), iterations x3 (10, 100, 1000)).

Psychedelic VHs are most commonly reported as being simple in nature (Kometer et al. 2011; Studerus et al. 2011; Schmid et al. 2015). These simple VHs may vary subtly in appearance dependent on the classical hallucinogen ingested (LSD, psilocybin, or DMT). In an attempt to capture variations in simple hallucinatory content, we used the bottom two layers of the DCNN (conv3 / conv4) to generate simple synthetic VHs. This led to an image set of 384 images (input images x32, AM type x2 (DGN-AM or DCNN-AM), DCNN layer x2 (conv3 or conv4), iterations x3 (10, 100, 1000)).

The experiment was hosted via the online experimental platform Gorilla (https://gorilla.sc/) and began by verifying that participants were eligible to take part in the study based on the pre-registered selection and exclusion criteria. Participants were asked to select a particular psychedelic experience from within the last 12 months, that they would use to answer all the questions in the survey. They were then asked to state the type of classical hallucinogen this experience related to, and to report the subjective potency of this experience using the following question: ‘How potent or powerful was your chosen psychedelic experience, in comparison to the most intense psychedelic experience you have ever had in your life? Please answer by giving a potency rating for your chosen psychedelic experience, on the following scale’, using a scale from 0 (not potent at all) to 5 (as potent as my most intense psychedelic experience).

Participants then completed a practice session before the main image selection task. The practice session was the same for all participants and consisted of 5 trials (3 complex and 2 simple), the images presented in these trials were randomly selected from the pool of images used in the main experiment. In each trial participants were presented with an array of 6 synthetic VHs (3 neurological and 3 psychedelic) and were asked to select, using a mouse, the image that most closely resembled their chosen psychedelic experience (Figure S8).

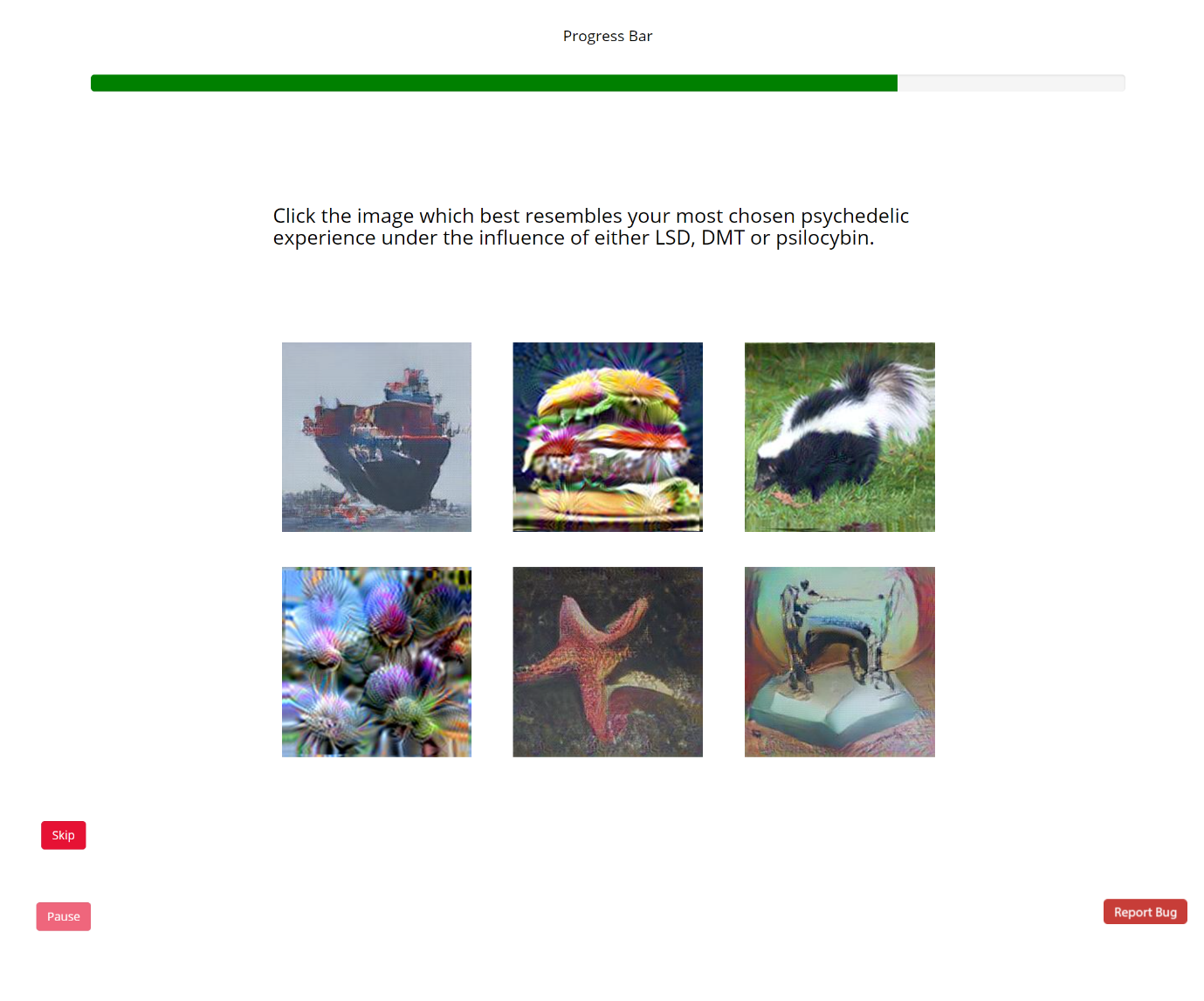

**Figure S8.** Sample trial presented to participants with recent psychedelic experience, in which they were asked to select the image that best resembled their most recent **complex** psychedelic experience. The images in each trial consisted of 3 complex psychedelic and 3 complex neurological synthetic VHs.

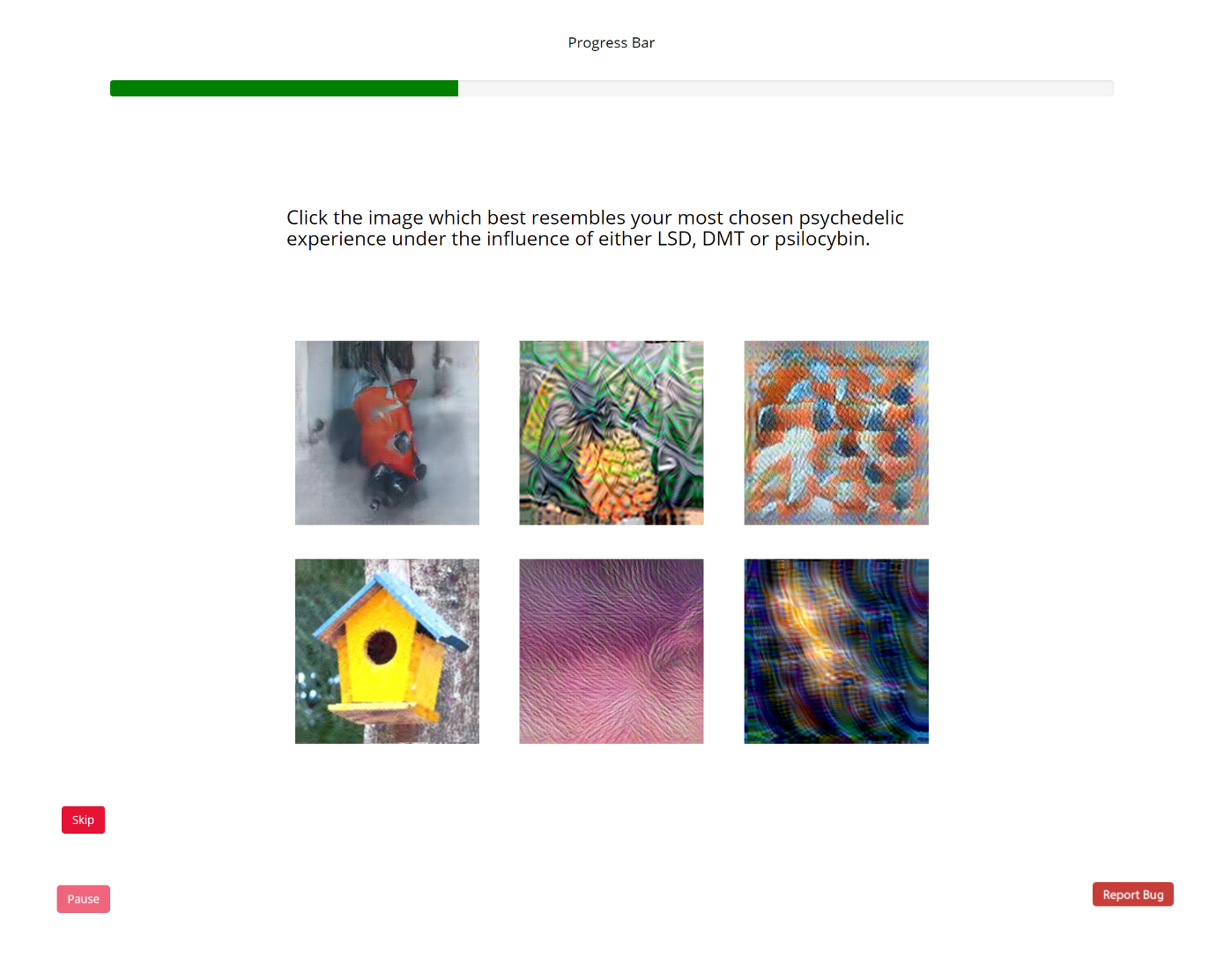

**Figure S9.** Sample trial presented to participants with recent psychedelic experience, in which they were asked to select the image that best resembled their most recent **simple** psychedelic experience. The images in each trial consisted of 3 simple psychedelic and 3 simple neurological synthetic VHs.

Participants had to complete each trial within 60 seconds and there was a 700 ms inter-trial interval. If none of the 6 images presented within a given trial matched a participant’s chosen psychedelic experience, they were instructed to press a ‘skip’ button to proceed to the next trial.

The main experiment used the same trial structure as the practice session but consisted of a total of four blocks, each with 32 trials. Two blocks contained images simulating simple VHs (3 CBS and 3 psychedelic), and two blocks contained images simulating complex VHs (3 neurological and 3 psychedelic). Block order was counterbalanced in an ‘ABBA’ manner for each participant. Each image was presented only once per block but may have been presented twice within the whole experiment. Image order was randomised for each participant and block. Within each block, 2 catch trials were placed in random positions. Catch trials contained a task irrelevant instruction (e.g., click the top-right image, or select the image that contains flamingos), and were included to maintain participants’ attention and prevent them from selecting images at random. Catch trials were also used to exclude participants from further analysis.

A chosen psychedelic experience may have contained both simple and complex VHs. To capture this information, we first provided participants with definitions of simple and complex hallucinations (see supplemental material for definitions) and then asked them to indicate if their chosen psychedelic experience involved simple, complex or both types of hallucinatory phenomena. Based on their response to this question they were asked to first report on the veridicality and then the spontaneity of their chosen psychedelic experience for either simple, complex, or both types of VHs. Participants were first provided with definitions of what was meant by the veridicality and spontaneity of a hallucinatory experience. Next, participants were asked to rate the veridicality of their chosen psychedelic experience compared to their normal visual experiences, on a scale from 1 (identical) to 10 (completely different). Participants were then asked separately about the spontaneity of their hallucinatory experience in relation to their existing visual experience on a scale from 1 (completely dependent) to 5 (completely independent). If participants chosen psychedelic experience contained both simple and complex VHs, the survey asked them about their complex VHs first.

**Questions used in psychedelic survey**

The first characteristic of your past psychedelic experience we want to find out more about is its complexity. Visual hallucinations can be categorised as simple or complex: Simple (or ‘non-formed’) hallucinations are hallucinations which are made up of extremely basic features. They often consist of small, bright spots, lights, colours, geometric shapes, or patterns. Complex (or ‘formed’) hallucinations tend to be clear, identifiable images or scenes. These might include visualisations such as faces, figures of real or bizarre humans or animals, objects, scenery, or entire landscapes.

Question 1: During your most recent psychedelic experience induced by LSD, DMT or Psilocybin, did you experience any complex visual hallucinations?

Response: [Yes/No]

Table S2. Question used in psychedelic survey to assess the complexity of chosen psychedelic experience.

On a scale from 1 (Identical) to 10 (Completely Different), how similar were your complex hallucination experiences (perception of identifiable forms: faces; objects; figures; landscapes; scenery) to your normal visual experiences, i.e., how these aspects appear when you have not ingested any hallucinogens?

Response: scale [1-10]

Table S3. Question used in psychedelic survey to assess the veridicality of chosen complex psychedelic VHs.

On a scale of 1 (Identical) to 10 (Completely Different), how similar were the simple aspects of this psychedelic experience (the lights, colours, lines, geometric shapes, structures and patterns) to the way these visual aspects appear when you have not ingested any hallucinogens?

Response: scale [1-10]

Table S4. Question used in psychedelic survey to assess the veridicality of chosen simple psychedelic VHs.

The second characteristic we want to find out more about is the dependency of your past psychedelic experiences. Visual hallucinations can be dependent upon or entirely independent from what you are seeing in the world:

Dependent visual hallucinations change or alter existing information within the visual scene, without adding any new content. This type of hallucination is a transformation of content already present in the visual field; thus, they are dependent upon what already exists. For example, seeing the brightness or colours of your environment fluctuate is a dependent hallucination.

Independent visual hallucinations do not alter what already exists in the visual field, but instead introduce entirely new spontaneous content. While this hallucinatory content may be consistent with the context and setting in which it appears, it is not a transformation of any existing content. For example, an image of a human or animal appearing superimposed at the center of your environment is an independent visual hallucination.

On the following scale from 1 (Completely Dependent) to 5 (Completely Independent), please indicate the extent to which the complex aspects of your psychedelic hallucinatory experience (perception of identifiable forms: faces; objects; figures; landscapes; scenery) were dependent upon or independent of existing visual content.

**Response:** scale [1-5]

Table S5. Description of spontaneity and question used to assess the spontaneity of chosen complex psychedelic VHs.

On the following scale from 1 (Completely Dependent) to 5 (Completely Independent), please indicate the extent to which the simple aspects of your psychedelic hallucinatory experience (the lights, colours, lines, geometric shapes, structures and patterns) were dependent upon or independent of existing visual content.

Response: scale [1-5]

Table S6. Question used to assess the spontaneity of chosen simple psychedelic VHs.

Note that participants were only answered questions about complex VHs if they indicated that their chosen psychedelic experience contained this type of hallucinatory content.

At the end of the experiment, participants were provided with a debriefing document, which explained the aims and hypotheses of the experiment. They were then informed if they had won an Amazon voucher or not. The experiment took approximately 30 minutes including instruction and debriefing, and the data was kept anonymous and confidential.

#### 1.3 Psychedelic Survey Supplemental Results

##### 1.3.1 ANOVA results examining the effects of the number of iterations and Error Function on image selection.

To investigate possible interactions between the image generation sub-categories and participants likelihood of selecting a synthetic VH in the image selection task, we conducted two separate ANOVAs for the psychedelic type, the type most frequently selected. The two ANOVAs were conducted on Error Function (Winner-Takes-All or Fixed) for complex VHs, and target layers (conv3 or conv4) for the simple VHs, and the number of iteration (10, 100, or 1000 iterations). For complex VHs the factors were (Iteration level (3), Error Function (2)) and for simple VHs (Iteration level (3), Layer of DCNN (2)). The 4 separate ANOVAs were: Complex-psychedelic, Complex-neurological, Simple-psychedelic, Simple-neurological.

**Complex- psychedelic**

No significant differences of iteration number or Error Function.

| **ANOVA - Selection Counts** | | | | | | | | | | |
| --- | --- | --- | --- | --- | --- | --- | --- | --- | --- | --- |
| **Cases** | | **Sum of Squares** | | **df** | | **Mean Square** | | **F** | | **p** |
| Iteration |  | 161.669 |  | 2 |  | 80.834 |  | 1.447 |  | 0.236 |
| Error Function |  | 5.194 |  | 1 |  | 5.194 |  | 0.093 |  | 0.761 |
| Iteration ✻ Error Function |  | 134.719 |  | 2 |  | 67.359 |  | 1.206 |  | 0.300 |
| Residuals |  | 22118.435 |  | 396 |  | 55.855 |  |  |  |  |
| *Note.*  Type III Sum of Squares | | | | | | | | | | |

**Simple- psychedelic**

The number of Iterations had a significant main effect while DCNN layer had no significant effect on image selection. Additional post-hoc tests (independent sample t-tests) revealed that iteration 10 was selected significantly more frequently than iteration 100 or 1000.

| **ANOVA - Response** | | | | | | | | | | | | |
| --- | --- | --- | --- | --- | --- | --- | --- | --- | --- | --- | --- | --- |
| **Cases** | | **Sum of Squares** | | **df** | | **Mean Square** | | **F** | | **p** | | **η²** |
| Iteration |  | 3170.161 |  | 2 |  | 1585.080 |  | 29.279 |  | < .001 |  | 0.123 |
| Layer |  | 113.533 |  | 1 |  | 113.533 |  | 2.097 |  | 0.148 |  | 0.004 |
| Iteration ✻ Layer |  | 44.303 |  | 2 |  | 22.152 |  | 0.409 |  | 0.664 |  | 0.002 |
| Residuals |  | 22520.711 |  | 416 |  | 54.136 |  |  |  |  |  |  |
| *Note.*  Type III Sum of Squares | | | | | | | | | | | | |

##### 1.3.2 Post Hoc Tests

| **Post Hoc Comparisons - Iteration** | | | | | | | | | | | | | | |
| --- | --- | --- | --- | --- | --- | --- | --- | --- | --- | --- | --- | --- | --- | --- |
|  | |  | | **Mean Difference** | | **SE** | | **t** | | **Cohen's d** | | **p _tukey_** | | **p _bonf_** |
| 10 |  | 100 |  | 5.900 |  | 0.865 |  | 6.824 |  | 0.774 |  | < .001 |  | < .001 |
|  |  | 1000 |  | 5.727 |  | 0.893 |  | 6.416 |  | 0.700 |  | < .001 |  | < .001 |
| 100 |  | 1000 |  | -0.174 |  | 0.878 |  | -0.198 |  | -0.028 |  | 0.979 |  | 1.000 |
| *Note.*  Cohen's d does not correct for multiple comparisons. | | | | | | | | | | | | | | |
| *Note.*  P-value adjusted for comparing a family of 3 | | | | | | | | | | | | | | |
| *Note.*  Results are averaged over the levels of: Layer | | | | | | | | | | | | | | |

#### 1.4 Clinical semi-structured phenomenological interview

Semi-structured interview used to enquire about the precise visual phenomenology associated with CBS and neurological VHs. All participants were interviewed by video conference call (zoom etc.) or telephone, which lasted no longer than 45 minutes.

Background info:

- Condition that has led to the experience of VHs:
- Current Medication:
- Age:
- Gender:
- Visual Acuity: Ask participant to describe the researcher’s appearance

Cognitive Ability:

- Note impression of Cognitive Ability from interview session

Background information of experience of visual hallucinations

- How long have you experienced VHs?
- What is your first memory of experiencing a VH?
- Visual Quality: Have your VHs changed in their quality over time? This could be in terms of their visual characteristics, colour, sharpness, detail or in terms of their content.
  - If not, for how long have they been consistent with today?
- Triggers: Do your VHs occur at a particular time of day, or are there particular triggers e.g., light level, tiredness?
- Frequency: As of today, what is the current frequency of your VHs? Has the frequency of your VHs varied over time?
- Reality monitoring – have you ever confused any of your VHs as being real?
  - If not, how are you aware that these experiences are not real?
- Spontaneity – if unclear from descriptions given below: Are your hallucinations transformations of things that you see in the world? Or are they independent of visual information that you see?

Interview questions:

1. When was your most recent VH?
2. Can you describe the content of this experience, where were you when it occurred?
3. Can you describe the colours associated with this experience? Normal colours, dim/bright colours
4. How life-like was the VH? Ask participant to look at and consider an object in their visual field, compared to looking at this object how similar are your VHs in visual quality.
5. How was it similar or different to your normal visual experiences is your experience of VHs, (allow participant to elaborate on the experience).
6. Could you give me as much detail as possible about the experience? Prompts: What size, distance, clarity etc., was the VH.
7. On a scale of 1-10, 10 being identical to normal visual experiences, 1 being not at all like normal visual experiences, how would you rate the visual quality of this VH?

out of 10.

8. Now I would like you to think of other examples of VHs that you remember particularly vividly. (Record notes for other examples as above).
9. Do any of your VHs relate to people you have known or items from your experience or memories?
10. Apart from the examples you have described, what other types of things do you hallucinate, e.g., people, animals, objects, insects.
11. If you experience hallucinations of people, can you describe what type of clothing they wear?
12. Now I would like you to think about different types of VHs you may have experienced, clinically we describe two main forms of VH, 1. Simple VH which are characterised by abstract shapes and repeating patterns such as grids. 2. Complex VHs are characterised by more identifiable forms such as faces, animals, objects, and scenes.
    a. Have you experienced Simple VHs?
    b. Could you give me a specific example of a Simple VH?
13. Next, I will show you a series of images, please select the column, if any, that displays the closest visual similarity to your experience of visual hallucinations.
